# A surface-engineered microfluidic device for antibody-mediated negative selection of high-quality sperm for assisted reproduction

**DOI:** 10.1101/2025.09.02.673619

**Authors:** Soraya Rasi Ghaemi, David J. Sharkey, Nicole O. McPherson, Markos Negash Alemie, Krasimir Vasilev, Sarah A. Robertson

**Affiliations:** Research Associate, Robinson Research Institute and School of Biomedicine, The University of Adelaide, South Australia 5005, Australia; Senior Research Associate, Robinson Research Institute and School of Biomedicine, The University of Adelaide, South Australia 5005, Australia; ARC Research Fellow (DECRA), Robinson Research Institute, Freemasons Centre for Male Health and Wellbeing and School of Biomedicine, The University of Adelaide, South Australia 5005, Australia, and Genea, Sydney, New South Wales; Biomedical Nanoengineering Laboratory, College of Medicine and Public Health, Flinders University, South Australia 5042, Australia; Matthew Flinders Professor, Biomedical Nanoengineering Laboratory, College of Medicine and Public Health, Flinders University, South Australia 5042, Australia; NHMRC Investigator Fellow, Robinson Research Institute and School of Biomedicine, The University of Adelaide, South Australia 5005, Australia

**Keywords:** Plasma polymerisation, Polyoxazaline, Gold nanoparticles, Functionalized surface, Sperm, Antibody, DNA fragmentation, Assisted reproduction

## Abstract

Preparation of spermatozoa with optimal developmental competence remains a challenge in assisted reproduction. Conventional techniques based on sperm motility and morphology fail to adequately remove sperm with DNA damage. Here, we report development of a microfluidic device with a functionalized surface, inspired by the physiological processes of immune cell-mediated sperm selection in the female reproductive tract. A plasma-polymerized polyoxazoline (PPOx) film is applied to glass channel slides by deposition of 2-methyl-2-oxazoline, to establish a stable, biocompatible interface confirmed by X-ray Photoelectron Spectroscopy (XPS), ellipsometry, and sperm culture assays. To selectively eliminate pre-apoptotic and apoptotic spermatozoa wherein DNA damage is common, anti-phosphatidylserine (Anti-PS) antibody is immobilized to the PPOx-coated surface proximal to the channel slide inlet, while the sperm chemoattractant progesterone is adsorbed near the outlet. To optimise selective functionality, the surface topography is tailored by covalent immobilization of gold nanoparticles and addition of microchannels. Sperm recovered after processing whole liquified semen then consistently exhibit high motility and morphology, with <1% showing apoptosis-associated membrane damage or DNA fragmentation. Compared with conventional swim-up or other microfluidic approaches, the device yields sperm with improved quality, offering a simple one-step sperm selection strategy with potential for application in human and animal assisted reproduction.

**Short text and graphic for 45 the Table of Contents (ToC):** This study reports a microfluidic device with a functionalized surface utilizing a polyoxazoline coating and covalently immobilized gold nanoparticles and anti-phosphatidylserine antibody. The device selectively eliminates pre-apoptotic and apoptotic spermatozoa and yields sperm with substantially improved quality and low DNA damage, offering a simple one-step sperm selection device with potential for application in human and animal assisted reproduction.

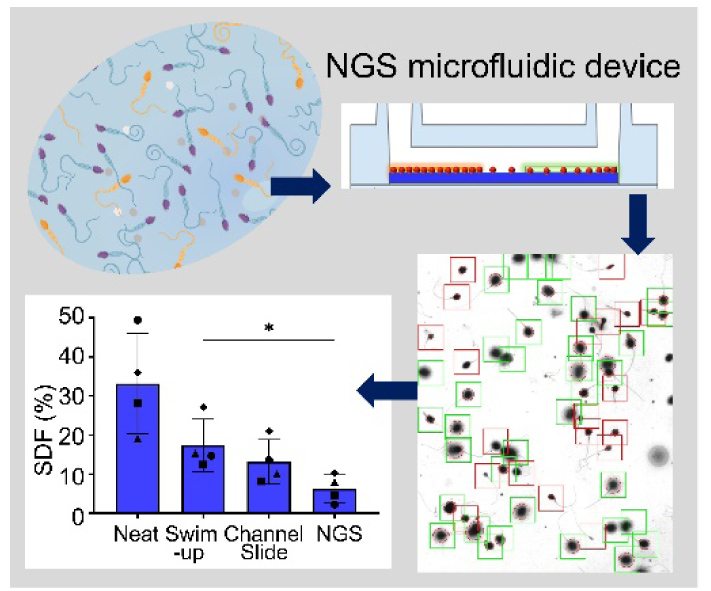

## 1. Introduction

Assisted reproductive technologies (ARTs) are widely utilized to treat human infertility and manage livestock production, but the success and efficiency of ART interventions are hampered by suboptimal protocols for preparation of sperm for oocyte fertilization and artificial insemination, and are inferior to the physiological processes occurring in the female reproductive tract.^[1]^ While ex vivo sperm selection has conventionally relied on physical parameters of sperm motility and morphology,^[1-2]^ these factors alone do not accurately predict a sperm cell’s ability to fertilize an oocyte and generate a healthy pregnancy, notably because DNA damage is common even in sperm with normal motility and morphology.^[3]^ This lack of precision in selecting spermatozoa means contemporary ART practice causes oocytes to regularly be fertilized using sperm with suboptimal DNA integrity and developmental competence, which can result in fertilization failure, poor embryo development, and later pregnancy loss.^[1b, 4]^ In response to these limitations, studies over the past decade have advanced techniques aiming to reliably generate high-quality spermatozoa for use in ART.^[2, 5]^ The Hyaluronic Acid (HA) Binding assay,^[6]^ Annexin V Magnetic Activated Cell Sorting (AV-MACS),^[7]^ ZyMot^®^ and related microfluidic systems,^[5a, 8]^ and the Zeta Method^[9]^ provide sperm with superior motility and morphology parameters - but each of these have practical shortcomings and whether any consistently translate to improved IVF outcomes is unclear.^[3c, 5b, 10]^

There is considerable potential for application of surface chemistry approaches in ART, including in the selection of high-quality sperm for *in vitro* fertilization (IVF). Surface modification to tailor interactions at the interface of materials has profound utility for understanding and recapitulating various biological processes.^[11]^ In cell biology, surface modification can be utilized to selectively regulate cell behaviour by modulating cellular adhesion, signaling, and communication.^[11b, 12]^ Often this is achieved through targeting the outer membrane, by which cells interact with surrounding biomolecules and extracellular matrixes, to in turn regulate many processes that impact cell function and fate.^[11b]^

Recent advances in modifying solid phase surfaces have paved the way for innovative approaches that leverage diversity in molecular structures on sperm cell membranes and their relationship to developmental potential. The outer membrane of spermatozoa plays critical roles in viability, metabolism, signaling, capacitation, acrosome reaction, and sperm-egg interaction.^[13]^ Due to its accessibility and close association with sperm quality, the sperm surface is a logical target for selecting high-quality spermatozoa.^[14]^ Of particular interest are surface molecules marking sperm in the early stages of apoptotic cell death that in the physiological setting of the female reproductive tract would destined for immune cell binding and sequestration,^[5b, 15]^ particularly as apoptosis is accompanied by DNA damage^[16]^. We reasoned that functionalized surfaces with specific ligands involved in immune sequestration could capture suboptimal sperm, thus mimicking the natural female reproductive tract environment, enabling preparation of sperm with improved parameters of developmental competence and fertility potential. Phosphatidyl serine (PS), which is confined to the inner leaflet of the plasma membrane in viable cells, becomes exposed during the early events of the apoptotic program of cell death.^[17]^ The presence of this surface marker commonly coincides with DNA damage.^[16]^ While sperm with externalized PS can remain motile, their compromised genetic integrity poses a higher risk of failure in fertilization or embryo development, or pregnancy loss.^[16]^

In vitro identification of pre-apoptotic sperm is challenging, as apoptotic changes are not detectable by morphological parameters alone – even under high-powered microscopy.^[18]^ Removal of sperm with externalized PS is the goal of the AV-MACS magnetic bead-based approach, but this technology involves exposing sperm to high powered magnetic fields and potential contamination of sperm cells with heavy metals.^[7a]^ In the current study, we aimed to develop an alternative method involving surface engineering that enables safe and efficient sequestering of apoptotic and pre-apoptotic sperm based on PS, thereby generating a residual pool of high quality negatively selected spermatozoa for recovery and use in oocyte fertilization. Here, we tailored the surface chemistry and topography of glass channel slides to achieve this goal. We utilized plasma polymerization of 2-methyl-2-oxazoline, resulting in the formation of a plasma-deposited polyoxazoline (PPOx) coatings able to facilitate the covalent conjugation of biomolecules targeting PS. We then explored surface topography modifications to alter roughness and geometry, aiming to promote sperm-biomolecule interaction. Gold nanoparticles (GNP) were incorporated to increase surface roughness, and microchannels were added to expand the reactive surface area and exploit the characteristic flagellar-propelled movements of sperm. The sex steroid hormone progesterone, which is known to exert chemotactic and maturational effects on sperm,^[19]^ was used to provide directional cues to facilitate sperm forward progression and promote developmental competence. Sperm quality after selection, compared with standard swim-up techniques, was assessed by measuring total motility, progressive motility, markers of early and late-stage apoptosis, and DNA fragmentation. The development of a functionalized surface platform for sperm selection has potential clinical applications in ART and opens the possibility of incorporating new biomolecules to facilitate spermatozoa preparation in the future.

## 2. Results and discussion

### 2.1. Plasma deposited PPOx film and characterization

PPOx films were deposited on glass microscope slides using continuous radio frequency power of 50 W for 2 minutes in the presence of 2-methyl-2-oxazoline precursor, in accordance with previous studies to optimize PPOx reactivity with biomolecules. ^[20]^ This process produced a hydrophilic PPOx film on the glass surface, characterized by a water contact angle of 56.3 ± 4.8 degrees. PPOx-treated surfaces allow covalent attachment of biomolecules by reacting with carboxylic acid (-COOH) groups to form irreversible amide bonds, ensuring stable biomolecular immobilization without risk of leaching from the surface substrate (Figure 1a,b).

**Figure 1.**
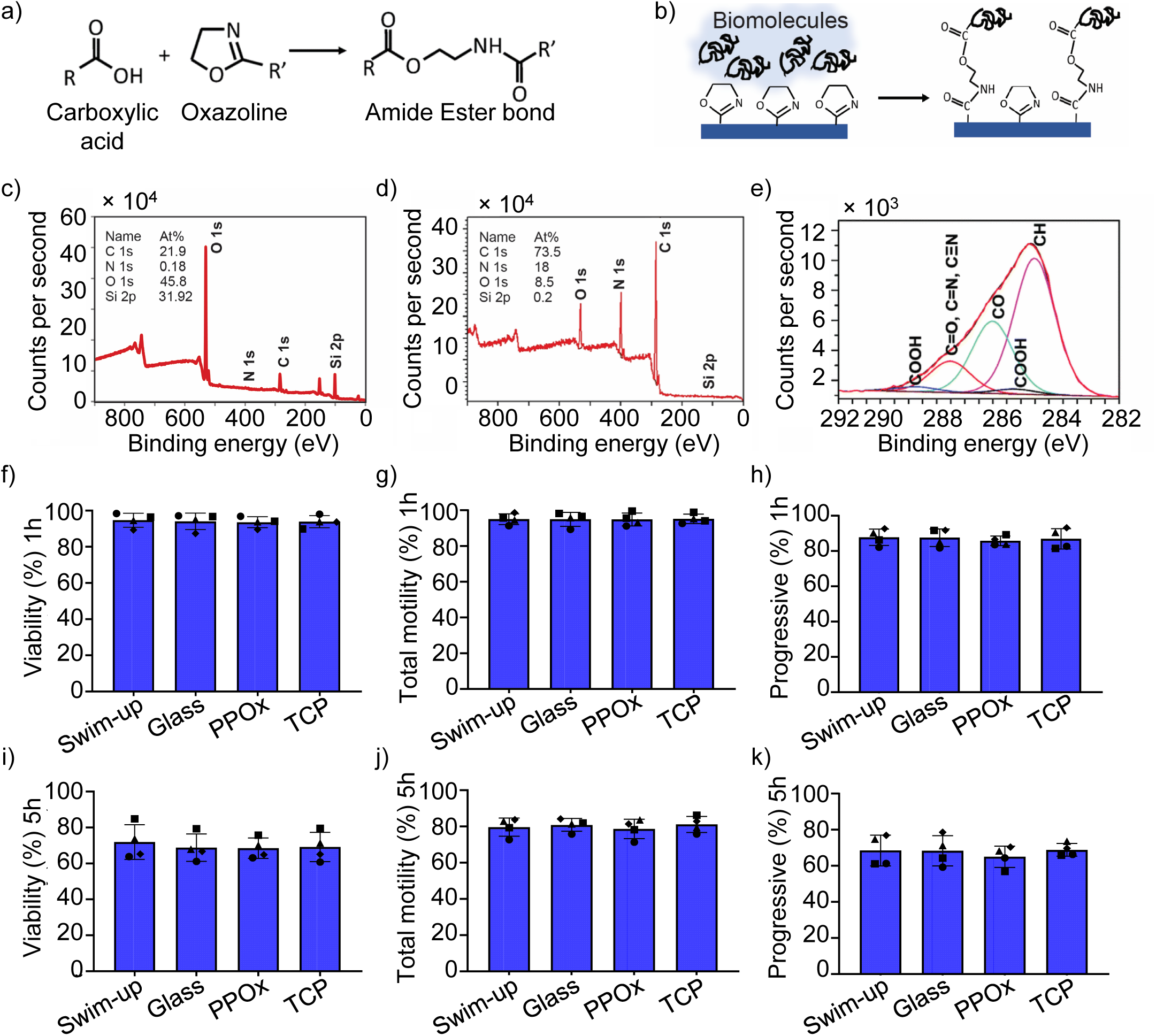
Surface modification by plasma polymerization and assessment of biocompatibility. (a) The chemical reaction between carboxylic acid and oxazoline. (b) Schematic of the bioconjugation interaction mechanism occurring on the PPOx-coated surface and carboxylic groups on biomolecules. (c) XPS spectra determining the atomic composition of the bare glass surface. (d) XPS spectra determining the atomic composition of the PPOx film. (e) High-resolution C1s spectrum (50 W, 5 min). (f-h) Biocompatibility assessment of PPOx-coated surfaces based on sperm viability, total motility, and progressive motility following 1-hour incubation with swim-up sperm on PPOx-coated coverslips (glass). (i-k) Analysis of biocompatibility of the PPOx-coated surface indicated by sperm viability, total motility, and progressive motility after 5 h incubation of swim-up sperm samples with PPOx-coated coverslips. All biocompatibility tests were performed at 37°C and 5% CO_2_. Different symbols show each of n=4 individual semen samples assessed.

The thickness of the PPOx films created via plasma deposition was measured using spectroscopic ellipsometry, revealing a thickness of approximately 26.1 ± 0.4 nm. The film’s durability and stability were tested by incubating coated slides for 24 hours with G-IVF^TM^ Plus, a clinical grade culture medium from Vitrolife. After incubation, the film’s thickness remained unchanged, indicating the PPOx coating was stable in a physiological environment.

When X-ray Photoelectron Spectroscopy (XPS) was used to analyze the chemical composition of the PPOx film, the atomic concentration of oxygen, carbon, and nitrogen was 8.5At%, 73At% and 18At%, respectively, consistent with the chemical structure of the precursor and published studies (Figure 1c,d).^[20b]^ The absence of a silicon signal confirmed the glass substrate was completely and uniformly covered by the PPOx coating, with no defects present (Figure 1d). The high-resolution C1s XPS spectra revealed that approximately 30% of the carbon atoms were in the C–O environment, and 10% in the C=O environment (Figure 1e). These reactive functionalities are consistent with the chemical structure of the oxazoline precursor as well as fragmentation and recombination reactions occurring in the plasma.^[20c]^ Notably, the spectra revealed signals specific to the oxazoline ring, confirming the presence of crucial C=N and C–O bonds.^[21]^

To evaluate the biocompatibility of the PPOx film, human sperm from normozoospermic donors prepared by a standard swim-up protocol were incubated on PPOx-coated glass coverslips in G-IVF^TM^ Plus medium for 1 hour at 37°C and 5% CO₂. Approximately 95% and 86% of spermatozoa retained total and progressive motility respectively on PPOx-coated coverslips, which was comparable with sperm incubated on bare uncoated glass coverslips or uncoated tissue culture polystyrene (Figure 1f–h). Similar data were obtained after 5 hours of incubation (Figure 1i-k), indicating that the PPOx coating did not negatively impact sperm motility and is biocompatible with human spermatozoa, as seen in other cell types.^[22]^

### 2.2. Surface characterization after biomolecule immobilization on PPOx coating

The unique reactivity of plasma-deposited PPOx with carboxylic acids has been exploited to irreversibly bind antibodies and other proteins to the plasma polymer substrate.^[23]^ This stable covalent attachment is particularly advantageous for clinical ART applications, as it ensures that immobilized molecules remain strongly attached to the substrate - minimizing the risk of their contamination of sperm preparations and contact with developing embryos. We next incubated Anti-PS monoclonal antibody at various concentrations with PPOx-coated glass coverslips and evaluated the degree of antibody binding using immunofluorescence staining after blocking unbound residues with 2% human serum albumin (HSA) (Figure 2a). Immobilized Anti-PS at concentrations of 50-400 μg/mL showed positive staining with FITC-labelled secondary antibodies, confirming a dose-dependent binding of Anti-PS onto the PPOx surface (Figure 2b-g). No staining was observed in the absence of Anti-PS antibody confirming the specificity of the reaction and the efficacy of HSA passivation (Figure 2c). Additionally, immunofluorescence staining confirmed that no delamination of the immobilized antibody occurred during incubation in G-IVF^TM^ Plus for 48–96 hours at 37°C.

**Figure 2.**
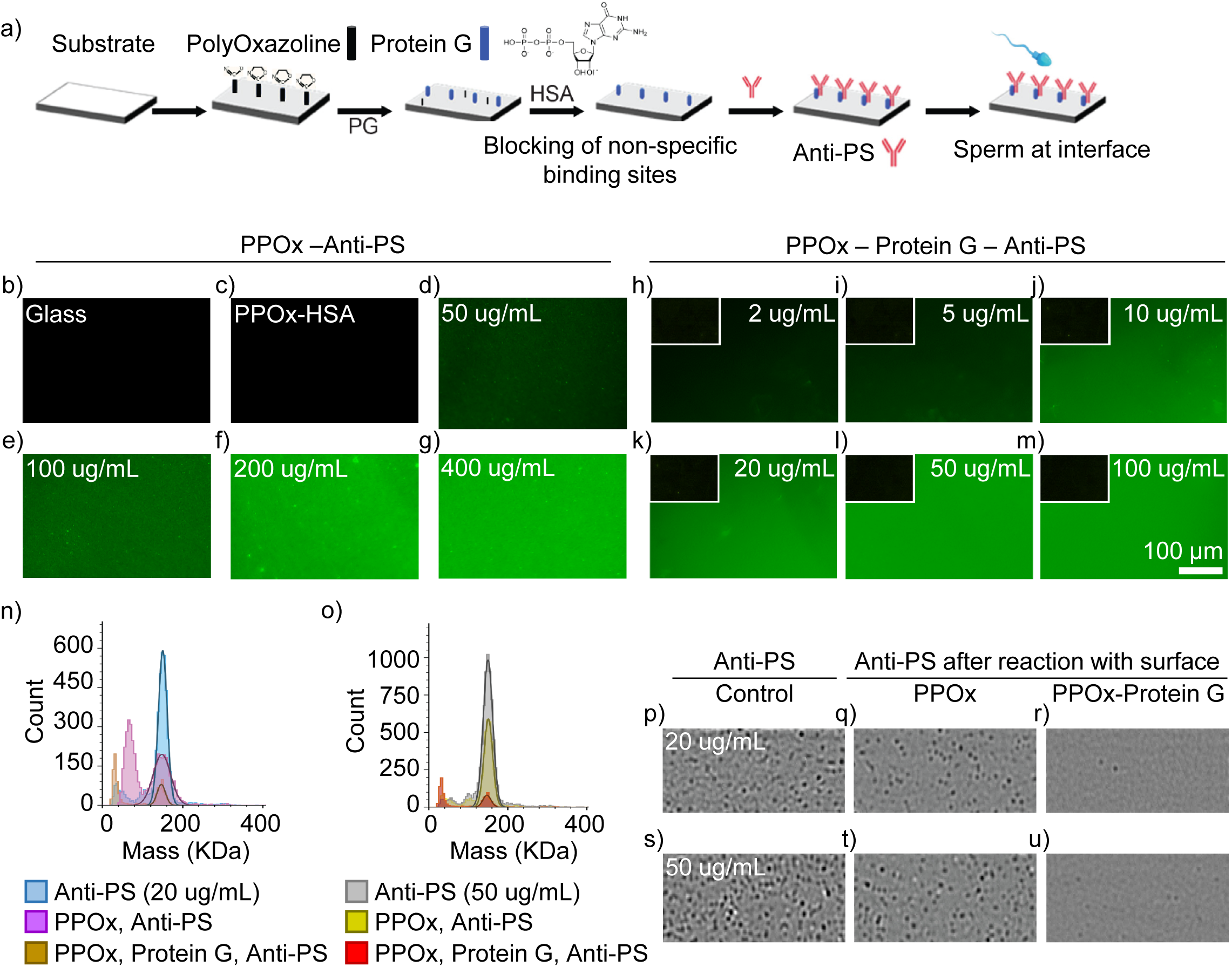
Surface immobilized anti-PS at different concentrations on PPOx-coated surface. (a) Schematic of Protein-G (PG) immobilization on the PPOx-coated surface and its interaction with anti-PS and spermatozoa. (b-g) Immunofluorescence staining of control and surfaces with immobilized anti-PS at concentrations of 50, 100, 200, and 400 μg/mL on PPOx-coated surfaces. After overnight antibody incubation at 4°C, surfaces were passivated with 2% HSA for 30-60 minutes. The presence of Anti-PS was revealed using FITC-labelled secondary antibody. Control surfaces treated with 2% HSA followed by FITC-labelled secondary antibody showed no binding. (h-m) Confirmation of protein-G on PPOx-coated surfaces using anti-Fn labelled with Alexa Fluor 488. PPOx-coated substrates were treated with or without protein-G, passivated with 2% HSA, and exposed to varying concentrations of anti-Fn labelled with Alexa Fluor 488, incubated overnight at 4°C. Fluorescence images show interactions at different anti-Fn concentrations (2, 5, 10, 20, 50, 100 µg/mL) with the protein-G PPOx-coated surface. Control reactions are in the upper left corner of each image. Scale bar: 100 µm. (n, o) Mass photometry assessment of protein-G function in immobilizing anti-PS on PPOx-coated glass. Relative density of anti-PS in solution (20 and 50 µg/mL) before and after interaction with PPOx-coated and PPOx-protein-G coated surfaces. (p-u) Mass photometer images showing anti-PS density in solution before and after incubation with different surfaces.

One of the key advantages of PPOx coatings is their ability to covalently immobilize biomolecules, which confers significant advantages in biomedical applications. Unlike surface chemistries like epoxy and aldehyde that bind amine groups,^[12a, 24]^ the PPOx coatings react with the carboxylic acid groups of proteins in a manner demonstrated to preserve protein bioactivity.^[20c]^ This binding mechanism is successful for a variety of proteins, including streptavidin, fibronectin, and anti-human podocalyxin antibodies, demonstrating its versatility.^[20b, 20d, 22a, 25]^

We next investigated the utility of incorporating protein G, which forms stable amide bonds with PPOx,^[11b, 12a, 20c]^ in order to allow a higher density of antibody absorption to the surface in a uniformly oriented manner that might afford increased efficacy for sperm sequestration.^[26]^ Protein G binds the Fc region of immunoglobulin proteins and so ensures precise antibody alignment on surfaces, which can be vital for accurate biosensor applications^[26b]^ and improves functionality in nanomedicine.^[27]^ The PPOx surface was incubated overnight with protein G prior to antibody incubation (Figure 2a). An indirect immunofluorescence approach using anti-fibronectin monoclonal antibody (Anti-Fn-Alexa 488) was then used to analyze protein G-mediated antibody capture, as direct methods using XPS were not possible due to the similar atomic composition of PPOx and biomolecules.^[20c]^ The PPOx-coated surface was incubated with protein G overnight at 4°C, then blocked with 2% HSA prior to application of Anti-Fn-Alexa 488 at concentrations of 2-100 µg/mL overnight. After washing with PBS, fluorescence microscopy revealed concentration-dependent binding of Anti-Fn-Alexa 488 (Figure 2h-m). Bare glass incubated with protein G, and PPOx-coated surfaces back-filled with 2% HSA without protein G, showed negligible fluorescence, indicating minimal binding of antibody without PPOx coating or after HSA blocking.

Mass photometry was also used to assess the utility of protein G. After overnight incubation of PPOx and protein G-PPOx coated surfaces with anti-PS at concentrations of 20-200 µg/mL, the residual antibody solutions were analyzed for remaining relative antibody concentration. Anti-PS antibodies bound to both surfaces, but with notably higher absorption to protein G-coated PPOx, indicated by a lower relative concentration of remaining antibody in solution in the latter case (Figure 2n-u). These findings add to other studies showing the utility of protein G in functionalized surfaces requiring antibody binding. For example, enhanced antibody binding efficiency through protein G has been shown to improve the detection of pathogens like influenza by affording lower detection thresholds.^[28]^ Similarly, Ryu et al. demonstrated that protein G increased sensitivity in lateral flow assays for detecting cardiac troponin I by ensuring specific antibody orientation on magnetic beads.^[26b]^

### 2.3. Sperm interaction with Anti-PS and Annexin V-coated surfaces

Next, we investigated whether anti-PS and Annexin V immobilized on glass coverslips can selectively sequester subpopulations of sperm. The aim was to assess their efficacy in capturing early-stage apoptotic spermatozoa, segregating them from non-apoptotic cells. Anti-PS and Annexin V were chosen as agents that mimic neutrophil interactions with sperm in the female reproductive tract.^[1b, 29]^ The effect on proportion of apoptotic sperm was evaluated by flow cytometry, using Annexin V as a marker of externalized PS indicating an early apoptotic state, and propidium iodide (PI) uptake indicating late stage apoptosis / cell death.

To assess the impact of Anti-PS antibody concentration on sperm attachment to coated surfaces, 1 × 10^6^ sperm from normozoospermic donors prepared by swim-up were added for 1 hour to coverslips coated with varying concentrations of immobilized anti-PS (50-400 µg/mL), initially without protein G. Spermatozoa attachment was visualized by light microscopy (Figure 3a-d), and bound sperm were quantified using a bight field microscope (Figure 3e). Anti-PS concentration significantly affected the proportion of sperm attaching, with maximal attachment observed at 200 and 400 µg/mL (Figure 3e).

**Figure 3.**
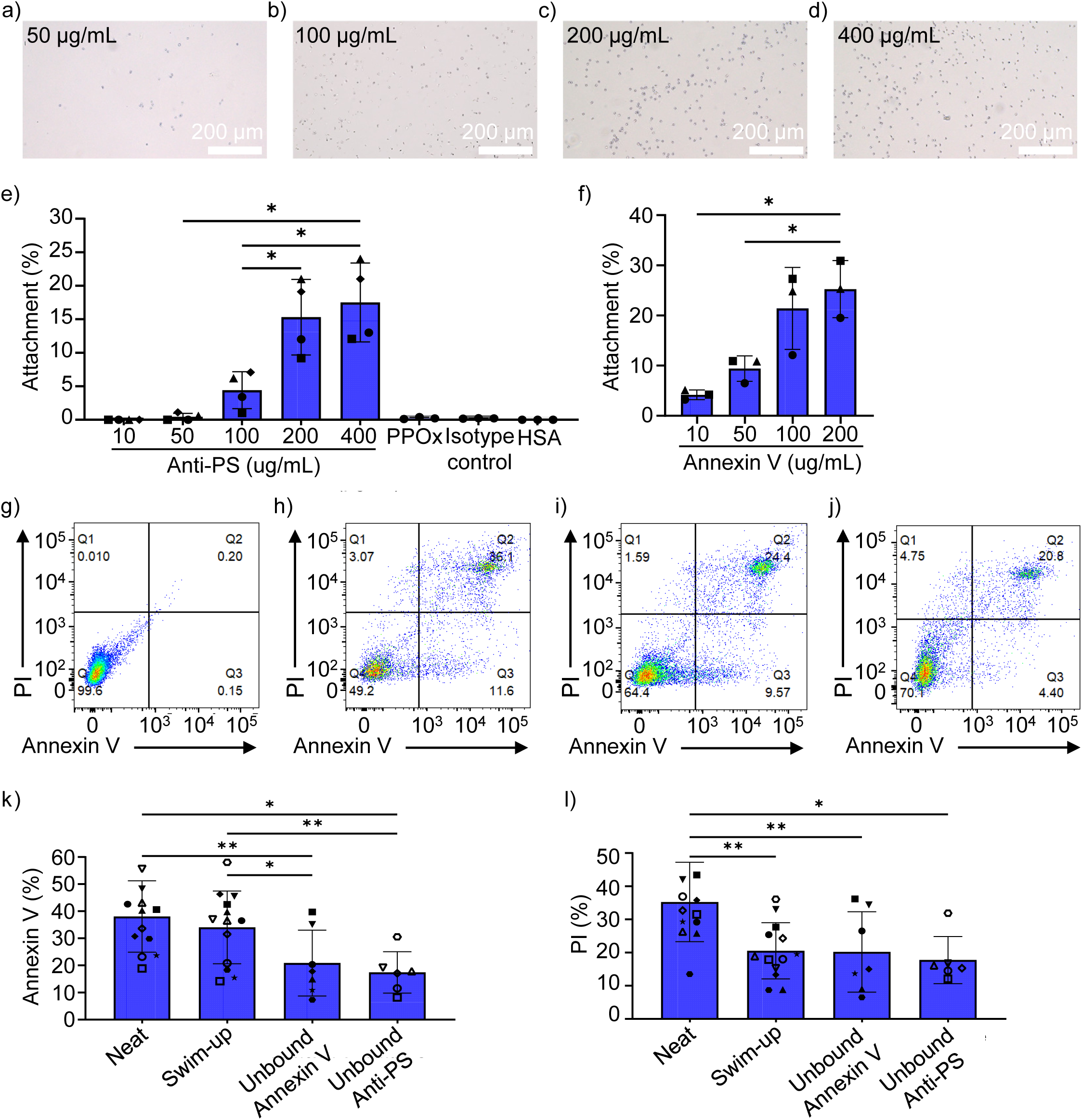
Sperm selection through incubation of swim-up sperm on glass coverslips functionalized with Annexin V and anti-PS. (a-d) Representative light microscope images demonstrating the attachment of swim-up sperm to the surface immobilized with anti-PS antibody at increasing concentrations (50, 100, 200, and 400 µg/mL) respectively. (e) The proportion of swim-up sperm at a density of 1 × 10^6^ in G-IVF^TM^ Plus adhered to the surface immobilized with anti-PS at various concentrations after 45-60 minutes of incubation. (f) The proportion of swim-up sperm at a density of 1 × 10^6^ in G-IVF^TM^ Plus adhered to the surface immobilized with Annexin V at various concentrations after 45-60 minutes of incubation. (g) Analysis of sperm apoptosis: a representative flow cytometry plot depicting gating strategy. (h) A representative flow cytometry plot depicting the neat sample. (i) A representative flow cytometry plot illustrating the swim-up sample. (j) A representative flow cytometry plot of unattached sperm after incubation on surface-immobilized anti-PS (200 µg/mL) coverslips. (k) The proportion of sperm labeled with Annexin V was quantified by flow cytometry analysis. (l) The proportion of necrotic or dead sperm was assessed using PI staining via flow cytometry. One-way ANOVA followed by post-hoc t-test was used to evaluate differences between treatment groups. *P ≤ 0.01 and **P ≤ 0.001.

The proportion of sperm attached to coverslips coated with 200 µg/mL Anti-PS was evaluated using sperm samples from six individuals to investigate between-individual differences. The proportion of sperm attached to the Anti-PS-coated surfaces ranged from 12% to 22% (Supplementary Figure 1a). Sperm attachment was dependent on Anti-PS-mediated binding, as negligible binding was observed on surfaces coated with PPOx alone, 2% HSA, or an irrelevant isotype-matched control antibody (Figure 3e).

The efficacy of PPOx-treated coverslips with surface-immobilized Annexin V at different concentrations (10-200 µg/mL) was also assessed, as an alternative to Anti-PS. A concentration-dependent response was seen, with greatest sperm attachment at 100 and 200 µg/mL of Annexin V (Figure 3f). In samples from six individuals, sperm attachment to coverslips coated with 200 µg/mL of Annexin V ranged from 5% to 23%, with minimal variation across replicates (Supplementary Figure 1b). Similar degrees of sperm attachment were observed on Annexin V- and Anti-PS-coated surfaces, indicating comparable efficacy in mediating spermatozoa attachment.

To determine the effect of incubation with Anti-PS- and Annexin V-coated surfaces on sperm quality, unattached sperm were analyzed by flow cytometry to quantify Annexin V^+^ and PI^+^ sperm (Figure 3g-j). A significant reduction in Annexin V^+^ sperm was observed in unattached populations recovered from the Anti-PS-coated or Annexin V-coated surfaces, compared to neat semen and swim-up samples (P ≤ 0.01, Figure 3k). Fewer PI^+^ sperm were present after incubation with the Anti-PS-coated and Annexin V-coated surfaces compared with neat semen, similar to swim-up samples.

These quality parameters were comparable to those reported by others for sperm recovered after incubation with surface-immobilized Annexin V. In studies using Annexin V-conjugated magnetic microbeads (AV-MACS),^[18, 30]^ sperm were passed through a high-power static magnetic field to generate two populations, one eluted from the magnetic field and enriched for Annexin V^-^ sperm and one retained by the magnetic field and enriched for Annexin V^+^ positive sperm. The population enriched for Annexin V^-^ sperm demonstrated higher motility, reduced caspase 3 activation, better mitochondrial membrane integrity, and less DNA fragmentation than the retained sperm. Studies evaluating the AV-MACS approach demonstrated that combining density gradient centrifugation and magnetic bead-based sperm sorting is superior to other preparation methods in terms of selecting motile, viable, and non-apoptotic spermatozoa.^[31]^Additionally, use of AV-MACS sperm preparation in IVF may improve pregnancy and cleavage rates, and increase the number of 8-cell (day 3) embryos with non-fragmented blastomeres per oocyte compared to density gradient preparation methods with oligoasthenozoospermic men.^[32]^ MACS-based sperm preparation may allow higher clinical pregnancy rates compared to sperm preparation by PICSI^®^ (Physiological Intracytoplasmic Sperm Injection) dish.^[30]^A systematic review and meta-analysis of prospective randomized trials found the use of AV-MACS for sperm selection increased pregnancy rates compared to density gradient and swim-up techniques, although no difference in implantation or miscarriage rates was seen. AV-MACS has also been successfully used after sperm cryopreservation for achieving successful pregnancy outcomes.^[33]^ A limitation of the AV-MACS approach is that it involves exposure to static magnetic fields which can have detrimental biological effects, including inducing pro-inflammatory changes and reactive oxygen species (ROS) generation.^[34]^ These effects might negatively impact spermatozoa integrity and potentially increase miscarriage risk.^[35]^Additionally, concerns exist about the potential for iron microbead contamination of sperm preparations used for clinical IVF, as residual microbeads might be transmitted to the oocyte during fertilization. These concerns raise practical challenges for the application of MACS in sperm selection for IVF and prompted us to progress development of our alternative approach.

### 2.4. Optimization of a functionalized surface channel slide

To improve the selectivity of the functionalized surface, we next applied the PPOx-coating and Anti-PS to a commercially-sourced glass channel slide and incorporated a series of modifications to the surface topography and geography.

Initially we evaluated the effect of channel height (0.1, 0.2, 0.4, 0.6, and 0.8 mm) on sperm selection using semen obtained from normozoospermic donors. After liquefaction and channel preparation, 60-100 µL of semen was introduced into the inlet chamber, and incubation occurred for 45-60 minutes at 37°C with 5% CO_2_. Sperm were then retrieved from the outlet, and their motility and quality were assessed (n=3-8). No significant differences in relative yield were observed across different channel heights compared to swim-up (Figure 4a). A channel height of 0.4 mm was chosen for further experiments, as it allowed efficient liquid handling while minimizing reagent use.

**Figure 4.**
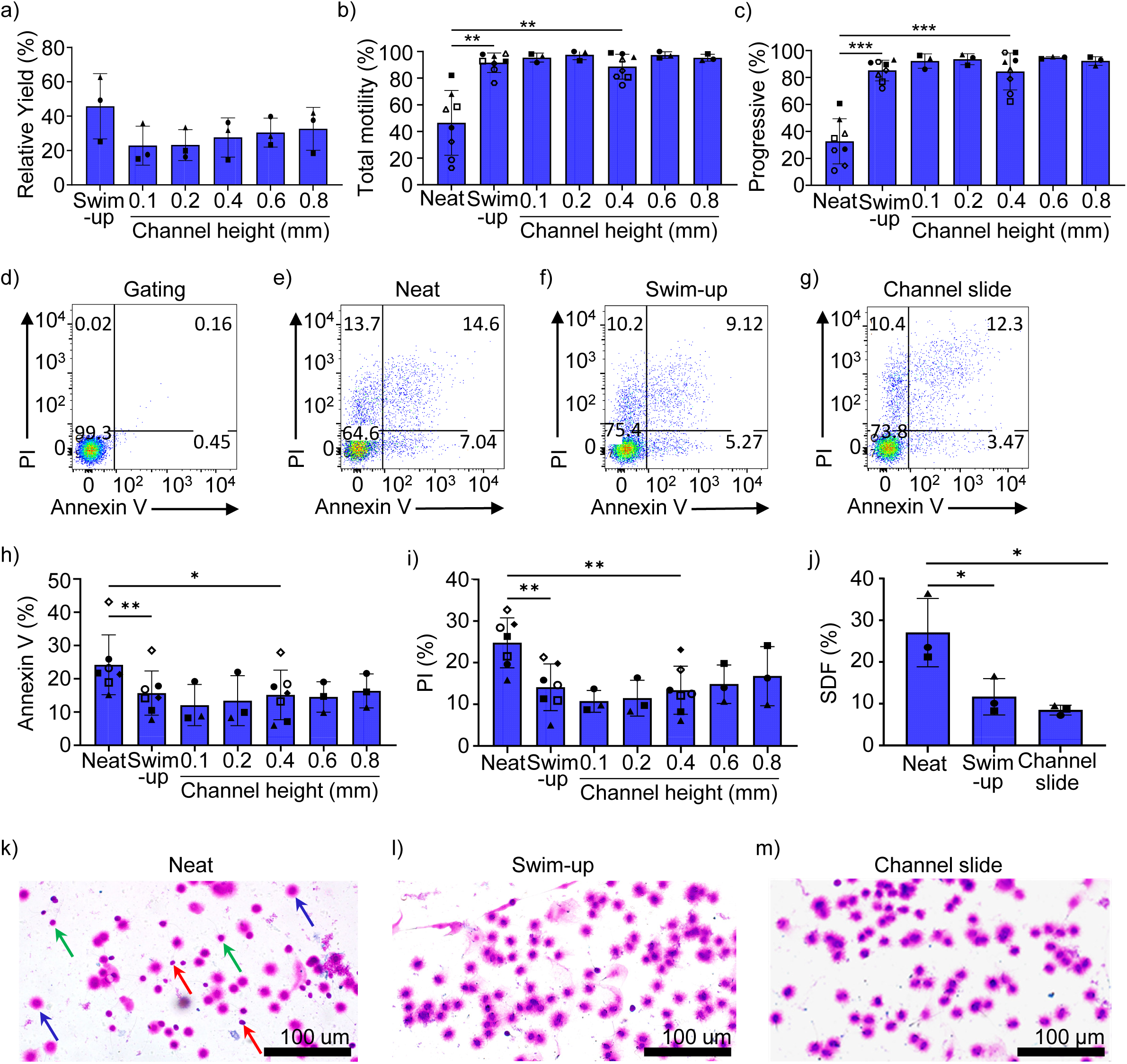
Evaluation of different channel heights on sperm recovery. (a) Comparison of the relative yield of progressive motile sperm obtained through swim-up and channel slide methods. (b, c) Total and progressive sperm motility assessed following collection via swim-up and channel slide methods. (d) Evaluation of the effect of different channel heights on sperm apoptosis: representative flow cytometry figures illustrating the gating strategy. (e) Representative flow cytometry figure of neat samples. (f) Representative flow cytometry figures of swim-up samples. (g) Representative flow cytometry figures showing sperm recovered from channels at a channel height of 0.4 mm. (h, i) Proportion of sperm labeled with Annexin V and PI, indicating apoptotic and necrotic populations, respectively, as evaluated by flow cytometry. (j) The proportion of sperm with DNA fragmentation. (k-m) Halosperm^®^ staining to determine the percentage of sperm population with DNA fragmentation in neat samples, sperm recovered through swim-up, and sperm recovered through channel slide, respectively. Red arrows indicate examples of DNA-damaged sperm with no halo. Green arrows indicate sperm with small halos and nuclei containing fragmented DNA. Blue arrows indicate sperm with large halos and nuclei with DNA integrity. One-way ANOVA followed by post-hoc t-test was used to evaluate differences between treatment groups. *P ≤ 0.01, **P ≤ 0.005, and ***P ≤ 0.001.

Spermatozoa recovered from the 0.4 mm channel slide showed high total motility and progressive motility comparable to swim-up, and significantly higher than neat samples (Figure 4b, c). The incidence of apoptosis measured by Annexin V staining in sperm recovered from channel slides were also comparable with swim-up and significantly lower than neat samples (Figure 4d-h), and the proportion of PI^+^ late stage apoptotic / dead sperm were similarly reduced (Figure 4i).

To assess the impact of channel slide and swim-up techniques on DNA integrity, we performed Halosperm^®^ staining on samples from three donors and determined the percentage of sperm with DNA fragmentation (SDF) (Figure 4j-m). Neat sperm samples showed an SDF rate of 27 ± 8%. Swim-up significantly reduced SDF to 12 ± 4%, while the channel slide method further decreased the SDF to 8 ± 1%. These results indicate that both methods effectively reduce SDF compared to neat samples.

These findings align with previous studies on microfluidic sperm selection. For instance, Shirota et al. demonstrated that microfluidic sperm sorters significantly reduced DNA fragmentation compared to conventional centrifugation and swim-up.^[36]^ Another study using a microchamber-based platform achieved efficient sperm separation with improved DNA integrity and higher motility.^[37]^

### 2.5. Optimization of Anti–PS concentration

This experiment investigated the effect of varying concentrations of immobilized Anti-PS on sperm selection using channel slides with a 0.4 mm height. Anti-PS was immobilized at concentrations of 50-300 µg/mL on PPOx-coated channel surfaces, and the effect on apoptotic sperm was evaluated using semen samples from three donors. After adding liquefied semen samples to the inlet chamber and incubating at 37°C and 5% CO₂ for 45 minutes, sperm recovered from the outlet were assessed for apoptosis incidence compared to sperm from neat and swim-up preparations. The Anti-PS coated-channel slide method significantly reduced Annexin V and PI positivity in sperm compared with neat samples (Supplementary Figure 2). The proportion of Annexin V^+^ and PI^+^ positive sperm was lowest at Anti-PS concentrations of 100 µg/mL or greater, showing effective reduction in the apoptotic population to a comparable or better degree than seen after swim-up, therefore we proceeded using 200 ug/ml Anti-PS.

### 2.6. Incorporation of gold nanoparticles (GNPs)

Next, we explored the impact on efficacy of sperm selection of increasing surface roughness by binding GNPs onto the coated surface. GNPs of various sizes (16, 38, and 68 nm) were incorporated after the initial PPOx coating, followed by a second ‘overcoat’ of PPOx, then Anti-PS coating. A reduction in Annexin V^+^ and PI^+^ sperm was seen in sperm recovered from GNP-coated channel slides compared with swim-up sperm (Supplementary Figure 3), that appeared to be an improvement over sperm recovered from smooth channel surfaces (Figure 4h, i). Comparable effects were seen for all GNP sizes, therefore we proceeded using 68 nm GNPs.

The improved quality of recovered sperm using GNPs aligns with studies showing that modification of surface topology and roughness can affect sperm-surface interactions, presumably by mimicking the features of membrane surfaces in the female reproductive tract.^[38]^ Rough surfaces, in particular, provide directional cues that enhance sperm adhesion and motility, and prevent clustering seen on smooth surfaces.^[3c, 39]^ The roughness of a surface increases its total contact area, allowing more interactions between the sperm cell membrane and the substrate.^[40]^ This can lead to better adhesion and interaction of sperm with the surface, promoting stable and directed movement.^[8]^ Smooth surfaces can cause sperm to accumulate in clusters due to the lack of directional cues, while rough surfaces can prevent this by providing pathways that encourage continuous movement.^[40-41]^

### 2.7. Incorporation of progesterone

While our channel slide method achieved comparable sperm quality to swim-up, the total recovery of progressively motile sperm was lower. We reasoned this limitation could be addressed by incorporating progesterone as a chemotactic factor to increase sperm directional movement. We next sought to identify an optimal concentration of progesterone for adsorption onto channel slide surfaces to support sperm function and viability. Progesterone at varying concentrations (1-100 µM) was adsorbed onto the surface near the outlet chamber of channel slides coated with PPOx and GNP (68 nm), then the remaining surface was backfilled with 2% HSA. Seminal fluid (60-100 µL) was introduced into the inlet chamber and incubated for 45 minutes. When recovered sperm from the outlet chamber were evaluated by Annexin V and PI staining, 5 µM and 10 µM progesterone were associated with the most favorable profiles (Supplementary Figure 4a, b).

To assess the release kinetics of adsorbed progesterone from functionalized surfaces, glass coverslips coated with carboxyl-functionalized GNPs were incubated with 10 µM of progesterone, then immersed in 150 µL of G-IVF™ culture medium at 37°C and 5% CO_2_ for various time points (0, 10, 20, 30, 40, 50, and 60 minutes). Media were collected and analyzed using a competitive ELISA. Progesterone quantification revealed a time-dependent release profile (n=3, Supplementary Figure 4c), with progressive release over 30 minutes and levels plateauing thereafter.

These results align with previous reports on the effect of progesterone on sperm function, and its ability to facilitate the biochemical and physiological changes that prepare sperm for fertilization.^[42]^ By interacting with receptors on the spermatozoon plasma membrane, progesterone induces calcium influx and activates signaling pathways essential for capacitation.^[42a]^ While progesterone is known to enhance sperm motility and viability,^[42b, 43]^ high concentrations are reported to induce oxidative stress, generating ROS that can damage sperm membranes and DNA^[44]^ and lead to sperm apoptosis.^[4a, 45]^ These considerations underscore the importance of having identified an optimal progesterone concentration.

### 2.8. Bio-conjugation of Anti-PS and progesterone with gold nanoparticles

We next developed a sequential coating strategy in the channel slide format (Figure 5). First, glass channel slides were coated with PPOx, followed by 68 nm GNPs. A second PPOx layer was then applied to the GNP-coated surface in the inlet chamber and half of the adjacent channel area, enabling covalent attachment of Anti-PS antibody (200 µg/mL). This partial overcoating process required masking the outlet chamber and adjacent half of the channel to preserve the carboxylic acid functionality on the GNP surface and enable subsequent non-covalent adsorption of progesterone (10 µM) onto the half of the channel adjacent to the outlet chamber. The resulting channel slides featured spatially distinct functional zones: one half covalently modified with Anti-PS to simulate the immune-mediated clearance of abnormal sperm, and the other half with non-covalently bound progesterone to enhance sperm chemotaxis and viability (Figure 5a).

**Figure 5.**
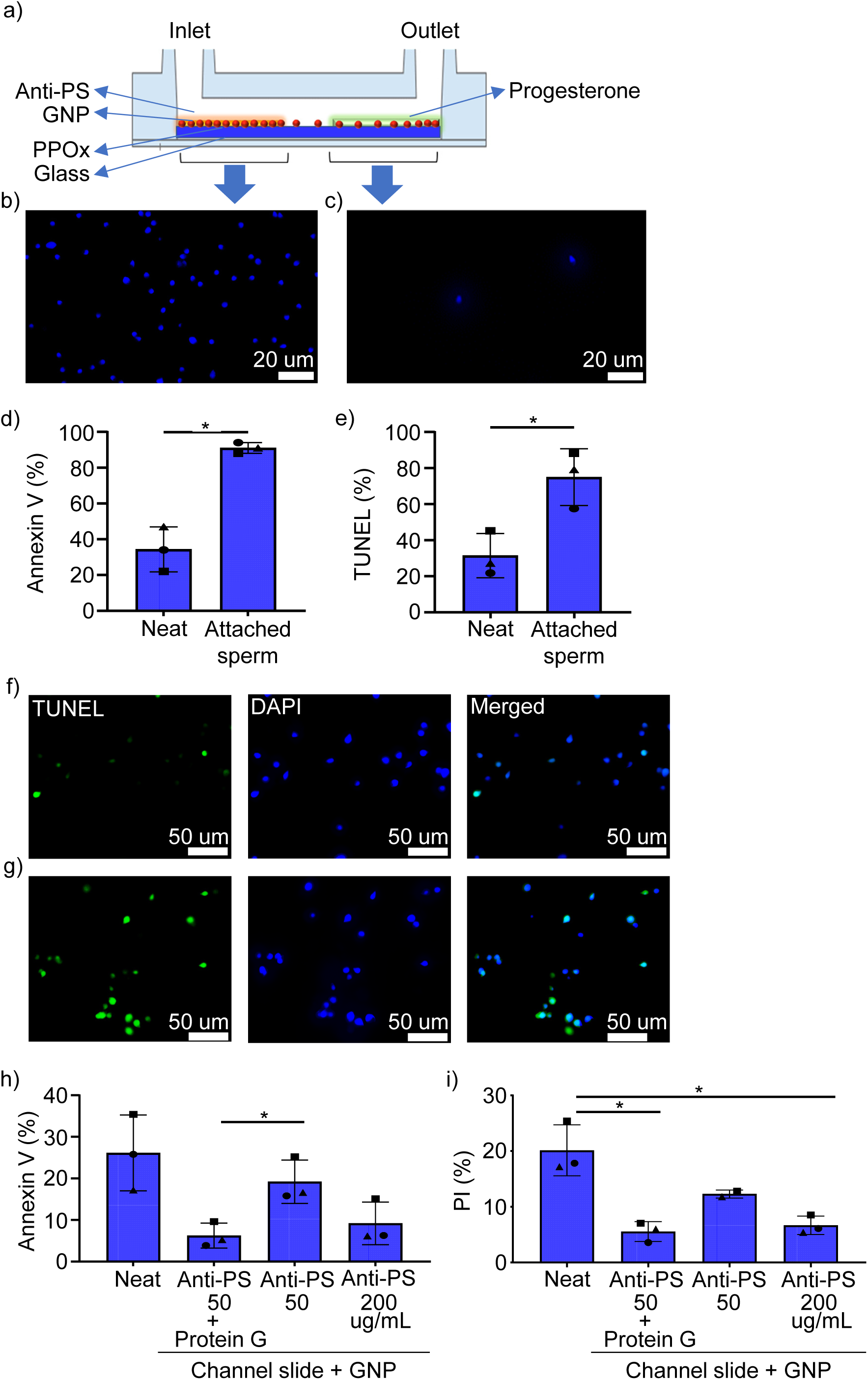
Fabrication of gold nano-particle-coated channel slides, containing areas of Anti-PS and progesterone coatings (the ‘NGS Device’) and its effect on immobilising low-quality sperm on the surface. (a) Surface coating with PPOx and addition of GNP to achieve covalent immobilization of COOH-GNP, followed by a second application of PPOx; covalent immobilization of anti-PS antibody (200 µg/mL) onto the glass surface next to the inlet well, and electrostatic adsorption of progesterone onto the half of the glass surface next to the outlet well. (b) Sperm attachment on the channel segment coated with Anti-PS. (c) The surface coated with progesterone exhibited a few or no sperm attachment. (d) Proportion of neat and attached sperm labelled by Annexin v was quantified by fluorescent microscopy. (e) The proportion of sperm labeled with TUNEL was quantified by fluorescent microscopy. (f) Representative fluorescent microscope images of neat samples stained with TUNEL (green) and DAI (blue). (g) Representative fluorescent microscope images of attached sperm on the neat and attached-sperm on Anti-PS-GNP coated surface stained with TUNEL (green) and DAPI (blue). (h, i) Bio-conjugation of Anti-PS using protein G and its effect on the reduction of apoptotic and necrotic sperm populations among recovered sperm through coated channel slides. One-way ANOVA followed by post-hoc t-test was used to evaluate differences between treatment groups. *P ≤ 0.01 and **P ≤ 0.001.

The effectiveness of this surface architecture in capturing and immobilizing spermatozoa was validated through fluorescence microscopy, with nuclear staining performed using DAPI (Figure 5b,c). After the removal of unattached sperm through washing, sperm attached to the surface were visualized using brightfield microscopy. A significant number of spermatozoa adhered to regions coated with anti-PS, particularly near the inlet chamber (Figure 5b). The density of sperm attachment decreased toward the progesterone-coated surface (Figure 5c). The proportion of apoptotic sperm and DNA fragmented spermatozoa immobilized on the surface was assessed by Annexin V and TUNEL staining, and capture of predominantly Annexin V^+^ and DNA fragmented sperm was demonstrated (Figure 5d-g).

We also evaluated the impact of incorporating protein G on the immobilization efficiency of the device. Utilizing a 0.4 mm channel slide coated with PPOx and 68 nm GNP, then overcoated with PPOx, we compared the selective efficacy of surfaces prepared by direct incubation of Anti-PS (50 or 200 µg/mL), or Anti-PS (50 µg/mL) immobilized using protein G to mediate its interaction with the PPOx surface. Sperm interaction was evaluated by comparing apoptotic and necrotic proportions among the unattached sperm populations (Figure 5h,i). Spermatozoa recovered from the surface coated with protein G-Anti-PS (50 µg/mL) demonstrated a significantly lower Annexin V^+^ population compared to surfaces coated with 50 µg/mL Anti-PS alone (P ≤ 0.01), and a comparable performance to that of surfaces coated with 200 µg/mL Anti-PS without protein G. The PI^+^ necrotic sperm population was significantly lower on protein G-Anti-PS compared to Anti-PS without protein G (P ≤ 0.01). These results underscore the utility of protein G in improving Anti-PS immobilization, allowing effective sperm binding at lower Anti-PS concentrations. These results are consistent with reports of utilizing protein G for directed immobilization of antibody via the Fc region,^[46]^ to prevent random orientation and thereby increase antibody density on surfaces.^[47]^ Previous studies have shown that use of Protein-G significantly enhances antibody affinity, potentially by up to 100-fold, compared to random antibody immobilization.^[46a]^ However, given that the attachment of protein G to antibodies is mediated non-covalently, there is a risk of unwanted detachment, potentially leading to contamination of spermatozoa. To avoid this issue we opted to continue device development without protein G, preferring to instead utilize a higher concentration of Anti-PS to maintain consistent binding efficacy and avoid the possibility of protein G contamination.

### 2.9. Efficacy of human sperm selection by the NGS Device

The preceding optimization steps culminated in the development of device we termed the nano-gold sperm (NGS) device designed to achieve selection of optimal quality sperm. In the next experiments we evaluated the selective efficacy of NGS devices comprising 0.4 mm channel slides coated with PPOx and 68 nm GNPs, then overcoated with PPOx before application of spatially distinct regions functionalized with Anti-PS (200 µg/mL) and progesterone (10 µM), and backfilled with 2% HSA. This configuration was evaluated for its ability to selectively isolate high-quality sperm from semen samples without any prior preparation.

Approximately 60-100 µL of neat liquefied seminal fluid was introduced into the inlet chamber of the NGS device, while the channel and outlet chambers were filled with sperm preparation medium. After incubation for 45 minutes, the recovered unattached sperm population were analyzed. Flow cytometry analysis (Figures 6a-e) showed the percentage of Annexin V^+^ sperm was lower in sperm recovered from NGS devices compared to neat sperm, sperm recovered from uncoated channel slides, or sperm prepared by standard swim-up technique (all P ≤ 0.05) (Figure 6f). A similar reduction was observed in the percentage of PI^+^ sperm.

**Figure 6.**
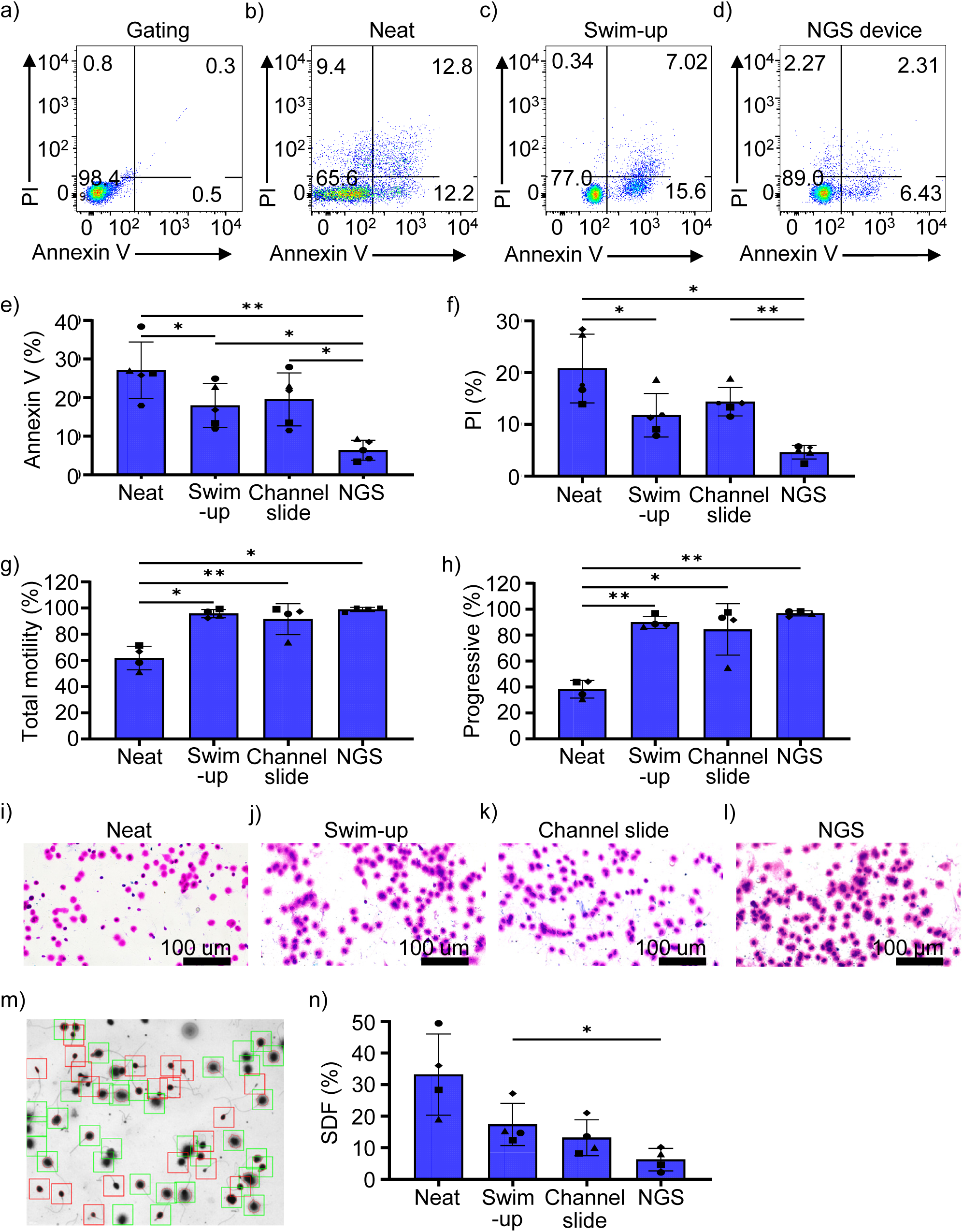
Comparison of the performance of the NGS device and conventional swim-up technique for sperm selection. (a) Illustration of the gating strategy in representative flow cytometry dot plots. (b) Flow cytometry dot plot representative of neat samples. (c) Flow cytometry dot plot representative of swim-up samples. (d) Flow cytometry dot plots depicting sperm recovered from the NGS device coated with Anti-PS at a concentration of 200 µg/mL without Protein-G. (e, f) The proportion of sperm labeled with Annexin V and PI, indicative of apoptotic and necrotic populations, respectively. (g, h) Analysis of sperm total motility and progressive motility following sperm selection using the NGS device or prepared by the swim-up technique, respectively. (i) Halosperm^®^ staining of neat samples of liquefied semen to determine sperm DNA fragmentation. (j) Halosperm^®^ staining of sperm after preparation using the standard swim-up technique to determine sperm DNA fragmentation. (k) Halosperm^®^ staining of sperm after preparation using a channel slide to determine sperm DNA fragmentation. (m) Halosperm^®^ staining of sperm after passage through the NGS device to determine sperm DNA fragmentation. (n) Analysis of halos via computer-assisted sperm analyzer (CASA), with green squares marking sperm with a medium to large halo, indicating unfragmented DNA, and red squares marking sperm with no halo or a small halo, indicating DNA fragmentation. (o) Quantification of the proportion of sperm with DNA fragmentation. Different symbols show each of n=4 individual semen samples assessed under each experimental condition. One-way ANOVA followed by post-hoc t-test was used to evaluate differences between treatment groups. *P ≤ 0.01, **P ≤ 0.005, and ***P ≤ 0.001.

Sperm motility was assessed following selection through the NGS device using CASA (Figure 6g). The results demonstrated a similar percentage of total and progressive motile sperm in samples recovered from the NGS device compared to the swim-up method, and improvement over neat sperm (P ≤ 0.001).

DNA fragmentation was assessed by Halosperm^®^ staining (Figure 6i) and CASA (Figure 6m). A significant reduction in the proportion of sperm with DNA fragmentation was observed after selection with the NGS device, compared with the swim-up technique and uncoated channel slides (Figure 6n).

These findings highlight the efficacy of the NGS device for selecting high-quality spermatozoa with low incidence of apoptosis, enhanced motility, and improved DNA integrity compared with the standard swim-up preparation technique, positioning it as a promising tool for application in assisted reproductive technology clinical settings.

### 2.10. Efficacy of human sperm selection by the NGS device with microchannels

To ensure robust enrichment of high-quality sperm with intact DNA integrity, we sought to further reduce sperm DNA fragmentation in sperm recovered from NGS devices. To achieve this, we further refined the device surface topography to more closely mimic the complexity of the female reproductive tract. Specifically, we introduced microchannels of a defined size onto the surface to increase the available surface area for sperm interaction and provide physical topographical cues that guide directional sperm movement. These microstructures mimic aspects of the natural architecture of the female reproductive tract, facilitating more physiologically relevant sperm navigation and potentially enhancing the selection of motile sperm. This modification was intended to accelerate the passage of high-quality sperm and minimize the passive drift of non-viable sperm, thereby decreasing the chance of dead or compromised sperm reaching the outlet chamber.

Microchannel structures were fabricated on a glass substrate (Figure 7a), onto which the afore-described surface modifications of the NGS device were subsequently transferred. These microchannels were precisely machined using a diamond saw under controlled conditions to ensure uniform depth and geometry. Following the fabrication process, profilometry analysis was conducted to characterize the microchannel dimensions. The results confirmed that the microchannels exhibited a depth ranging from 45–50 µm, a channel width slightly exceeding 100 µm, and an inter-channel distance of approximately 100 µm, with a predominantly U-shaped cross-sectional profile (Figure 7b). The total number of microchannels formed on each slide was 20–26.

**Figure 7.**
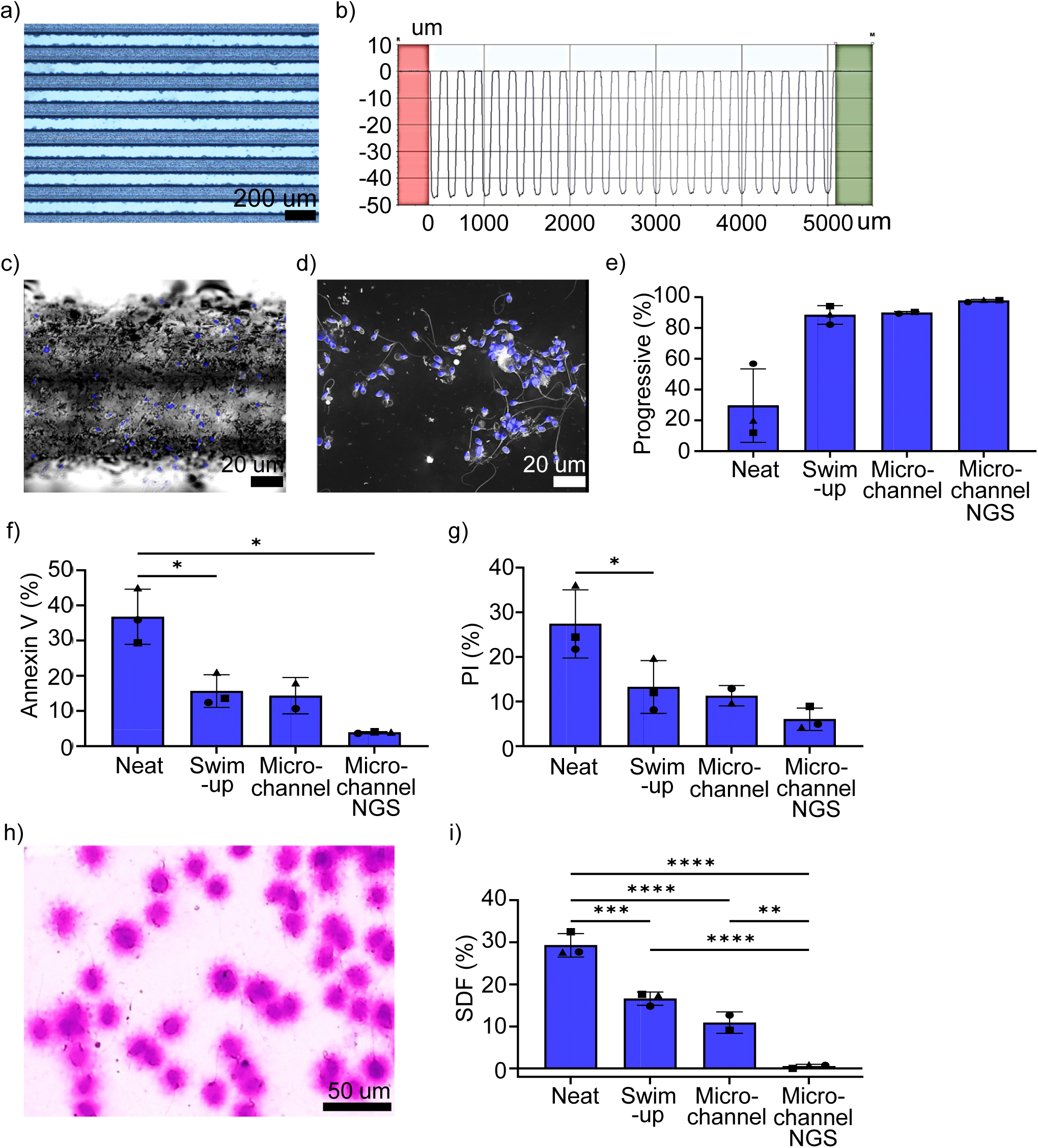
Fabrication of microchannel-NGS device. (a) Representative microscopic image of microchannel created by diamond saw on the normal glass slide. (b) Profilometry analysis of the microchannel dimensions. (c) sperm capture in the microchannel groove. (d) captured sperm between microchannel grooves. Nuclear stained by DAPI in blue and image was captured by holotomography microscope. (c) Analysis of sperm progressive motility following sperm selection using the microchannel-NGS device or prepared by the swim-up technique. (e, f) The proportion of sperm labeled with Annexin V and PI, indicative of apoptotic and necrotic populations, respectively. (g) Halosperm® staining of sperm after passage through the microchannel-NGS device to determine sperm DNA fragmentation. (h) Quantification of the proportion of sperm with DNA fragmentation. Different symbols show each of n=3 individual semen samples assessed under each experimental condition. One-way ANOVA followed by post-hoc t-test was used to evaluate differences between treatment groups. *P ≤ 0.01, **P ≤ 0.005, and ***P ≤ 0.001.

The functionalized microchannel-NGS device was employed for sperm selection using the previously validated protocol. A 100 µL aliquot of neat liquified seminal fluid was introduced into the inlet chamber, and spermatozoa were collected from the outlet chamber after an incubation period of 45 minutes. The attached sperm on the surface were imaged by Holotomography microscope (Tomocube HT-X1) (Figure 7c,d). The retrieved spermatozoa were analyzed for progressive motility, apoptotic status (Annexin V and PI staining), and DNA fragmentation (Halosperm^®^ assay). Selection efficiency was assessed by comparing key sperm quality parameters across four experimental groups: neat semen, swim-up, unfunctionalized microchannel slide (control), and microchannel-NGS.

Progressive motility improved significantly after sperm selection, with the microchannel-NGS system yielding the highest motility. Neat semen had the lowest motility, while swim-up and unfunctionalized microchannels showed similar efficacy (Figure 7e). This suggests that the microfluidic environment aids motile sperm selection, with the functionalized surfaces further enriching high-motility sperm.

The microchannel NGS device afforded a substantial further reduction in the proportion of Annexin V^+^ and PI^+^ sperm, compared to the device without microchannels (Figure 7f). The DNA fragmentation rate, assessed via Halosperm^®^ assay, demonstrated a further substantial reduction (Figure 7h-i), with a DNA fragmentation rate of 0.6 ± 0.5% (mean ± SEM), highlighting high efficacy in selecting sperm with intact genomic integrity.

These findings underscore the efficacy of the microchannel-NGS device in isolating sperm with optimal functional competence, surpassing conventional selection techniques in ART. A literature-based comparison of our microchannel-NGS system with established sperm selection approaches, including swim-up, density gradient centrifugation (DGC), Zeta potential selection, hyaluronic acid (HA) binding, AV-MACS, and microfluidic sorting (e.g., ZyMot, Fertile® chip), indicates superiority in all critical sperm quality parameters.^[2, 5a, 48]^ The proportion of Annexin V^+^ sperm, indicative of early apoptosis, was markedly lower in the microchannel-NGS system (3.9 ± 0.2%) compared with swim-up (15.7 ± 4.6%), density gradient centrifugation (∼10–25%), and HA binding (∼10–12%).^[49]^ AV-MACS which specifically targets apoptotic sperm removal, has a reported Annexin V^+^ sperm rate of 5– 10%.^[2, 7a, 30, 48c, 50]^ Although HA binding targets mature sperm,^[6b, 15]^ it does not prevent the selection of apoptotic sperm with fragmented DNA. AV-MACS and ZyMot offer reductions in apoptotic sperm, yet the NGS device yielded a more pronounced decrease. This suggests that the combination of surface functionalization and microfluidic-assisted selection effectively eliminate compromised spermatozoa while preserving the most viable population.

DNA fragmentation significantly impacts embryo quality and ART success rates. The microchannel-NGS system reduced sperm DNA fragmentation to 0.6 ± 0.5%, outperforming all conventional techniques. AV-MACS and HA binding yield sperm with reported DNA fragmentation rates of 8–13.5% and 2.5–15% respectively.^[51]^ Swim-up and density gradient centrifugation approaches have rates of ∼16-18% and ∼18–25% respectively, all of which are considerably higher than the rate seen in our microchannel-NGS approach.^[48b, 50, 52]^

Unlike AV-MACS, which relies on an antibody-based magnetic sorting mechanism,^[7a, 51a]^ or HA binding,^[51a, 51e]^ the microchannel-NGS system integrates surface topographical modifications and bioengineered functionalization to sequester suboptimal sperm. This dual-action approach, which mimics the natural sperm-filtering mechanisms of the female reproductive tract, allows a simple to use, minimally invasive, high-throughput selection of spermatozoa with superior motility, membrane integrity, and genomic stability. In contrast to conventional microfluidic sorting devices (ZyMot, FERTILE chip), which rely solely on hydrodynamic sperm migration,^[53]^ the microchannel-NGS system employs engineered surface properties to substantially improve its selective capability.

Given its superior motility enrichment, apoptotic sperm reduction, and DNA integrity preservation, the microchannel-NGS system represents a promising advancement for ART applications. By eliminating the need for centrifugation, magnetic field, or chemical-based selection techniques, this system may reduce oxidative stress and mechanical damage often associated with conventional and alternative methods. Future studies should explore its efficacy in improving fertilization rates, embryo development, and clinical pregnancy outcomes in ART. Additionally, further optimization of microchannel geometries, functionalization chemistries, and fluid dynamics may enhance its scalability for manufacture and broader clinical use.

## 3. Conclusion

In conclusion, this study demonstrates the utility of functionalized surfaces for ex vivo preparation of high-quality sperm, fabricated using plasma-deposited 2-methyl-2-oxazoline (PPOx) coating to enable covalent biomolecule application. Using an innovative approach of sequential coating with PPOx, gold nanoparticles, anti-PS antibody, and progesterone we created spatially distinct regions on glass channel slides to maximise spermatozoa-biomaterial surface interaction and optimize sequestration of suboptimal sperm. The utility of protein G to orient bound antibody and increase attachment of apoptotic spermatozoa was demonstrated, but was considered not necessary for robust performance.

The integration of microchannels into the surface modification further improved spermatozoa selection and notably benefited DNA integrity of recovered sperm. The microchannel-NGS device proved highly effective in isolating high-quality spermatozoa, as demonstrated by a marked reduction in apoptotic sperm and enhanced motility, as well as lower levels of DNA fragmentation in recovered sperm compared to conventional and alternative selection techniques. These findings demonstrate the potential of this functionalized surface device as a novel tool and superior process for enabling the selection of high-quality spermatozoa for assisted reproduction in humans and potentially in other species. Further evaluation of this tool in assisted reproductive technology clinical settings is now underway.

## 4. Materials and methods

### 4.1. Ethics approval and study participants

Approval for this study was obtained from the human research ethics committee of the University of Adelaide (H-2022-003) and all protocols followed The National Statement on Ethical Conduct in Human Research (2023). All participants provided written informed consent, were aged between 18 and 50 years, in good general health, and met WHO VI criteria for normozoospermia.^[54]^ Semen samples were produced by manual masturbation following a 2–7 day period of ejaculatory abstinence. Men with a history of vasectomy or vasectomy reversal, undescended testicles or genetic conditions affecting fertility (i.e. Prader-Willi, Klinefelter’s Syndrome), or blood-borne infections such as HIV/AIDS, were excluded.

### 4.2. Preparation of plasma deposited polyoxazoline thin films

Plasma polymerisation was achieved by deposition of 2–methyl–2–oxazoline (Sigma– Aldrich, Australia) onto oxygen plasma etched sticky–Slide I Luer (ibidi®) and standard glass slides in a capacitively–coupled bell chamber reactor, using established standard methods.^[20b, 20c]^ In brief, plasma deposition was performed using a plasma power of 50 W and 0.8 × 10^-1^ mbar pressure for deposition of oxazoline precursor for 2 min. The resultant plasma deposited polyoxazoline (PPOx) surface coating generates capacity to form irreversible covalent amide bonds through reacting with carboxylic acid groups.

X–ray photoelectron spectroscopy (XPS) (SPECS SAGE, Germany, monochromatic Mg Kα radiation source, 15 kV, 10 mA) was used to define the atomic composition of the PPOx film. Survey spectra were obtained at the pass energy of 40 eV with a resolution of 1 eV and measurement angle of 54° between the X–ray irradiation and the analyzer. Atomic concentrations were determined by analysis of the spectrum peak areas using Casa XPS software version 2.3.10 (Casa Software).

The wettability of the PPOx-coated glass slides was analyzed by sessile drop water contact angle measurements, with oxygen plasma-cleaned glass serving as the control. After surface polymerization, 5 μL Milli-Q water droplets (Eppendorf) were dispensed onto the surface using a motor-driven microliter syringe. Three points along the surface were evaluated in triplicate, and the process was monitored via video camera. Contact angles were then measured from images of the droplets using ImageJ software. The thickness of the PPOx films was determined using ellipsometry (Variable Angle Spectroscopic Ellipsometer, J.A. Woollam Co. Inc., USA), with data analysis conducted using WVASE32 software and the Drop Analysis plugin (NIH).

### 4.3. Gold nanoparticle (GNP) synthesis, immobilization, and characterisation

Gold nanoparticles (GNPs) were synthesized following a well-established protocol that reduces tetrachloroauric acid (HAuCl_4_, Sigma–Aldrich, Australia) with trisodium citrate (Na_3_Ct, Sigma–Aldrich, Australia) in an aqueous solution under a controlled reflux system.^[20a, 55]^ Specifically, a 0.01% HAuCl_4_ solution was heated to boiling while stirring, followed by the addition of Na_3_Ct as the reducing and stabilising agent, and the mixture was boiled for an additional 20 minutes. GNP size was controlled by varying the volume of 1% Na_3_Ct added, producing nanoparticles with diameters of 16, 38, and 68 nm. The GNPs were then capped with mercaptosuccinic acid (MSA, Sigma–Aldrich, Australia) to provide COOH surface functional groups.

For immobilization, COOH-functionalized GNPs were applied to PPOx-coated glass slides. The slides were immersed in GNP solutions overnight, followed by rinsing with Milli-Q water and drying under nitrogen gas. The slides were then again coated on one half with PPOx to facilitate covalent immobilization of anti-PS, while shielding the other half from PPOx to retain carboxylic acid groups for electrostatic adsorption of progesterone. GNP shape and size were characterized using transmission electron microscopy (TEM, JEOL– 2100F, Japan). Atomic force microscopy (AFM) was used to assess surface topography after GNP binding. Surface chemistry was analyzed using XPS survey spectra.

### 4.4. Microchannel formation on glass surfaces via diamond saw cutting

To create microchannels prior to the first PPOx coating, the glass substrate was cleaned and prepared to ensure a smooth surface free from contaminants. An automatic dicing saw (DAD3350, DISCO) was then used to create regular microchannels parallel to the long axis of the slide. The microchannels had a width of 100 µm and channel depth of 45-50 µm, dimensions optimized to minimize edge chipping.

### 4.5. Biomolecule immobilisation

Anti-PS antibody (clone 1H6, Catalog number: 05-719, Sigma Aldrich) and recombinant human Annexin V protein (Catalog number: NBP1-30265, In Vitro Technologies) were diluted in phosphate buffered saline (PBS, Thermo Fisher Scientific) to achieve concentrations between 10 and 400 µg/mL. Depending on the area to be coated, 20–70 μL of the biomolecule solution was carefully applied to the PPOx-coated glass substrate and incubated overnight at 4°C to allow binding. After incubation, the antibody solution was aspirated, and any unreacted PPOx groups were blocked with 2% human serum albumin (HSA, VitroLife) for 30-60 minutes at 4°C.

To enable orientation of the immobilized antibodies, protein G was used as an intermediary layer. A 1/10 dilution of protein G (Sigma-Aldrich, Merck) in PBS was applied to the activated PPOx surface and incubated overnight at 4°C. Following incubation, the surface was thoroughly rinsed with PBS to remove any unbound or loosely attached protein G molecules. The coated surface was then used to immobilize Anti-PS by the same procedure.

Before adsorbing progesterone onto the GNP-coated surface containing COOH groups, the surface was washed with PBS three times for 15 minutes each to eliminate unbound substances or GNPs. Subsequently, 0-100 µM of progesterone in a final volume of 30-50 µL was incubated with the surface overnight under desiccator. The interaction between progesterone and the COOH-coated surface is considered to involve reversible physical adsorption.

### 4.6. Surface-bound biomolecule characterization

Immunofluorescence analysis was performed to detect anti–PS antibody immobilized at different concentrations onto the PPOx-coated surface. After passivation with 2% HSA for 30-60 minutes the coated surfaces were washed with sterile PBS prior to use. Slides were then incubated with fluorescein isothiocyanate (FITC)–conjugated anti–mouse antibody (Abcam, diluted 1/500 in PBS for 30 minutes at room temperature). Incubation of secondary antibody with a PPOx-coated slide without prior anti–PS antibody coating served as a negative control.

To assess the relative concentration of Anti-PS after interacting with PPOx-coated and PPOx-PG-coated surfaces, mass photometry (Refeyn TwoMP, version 1.0) was employed. The data obtained from this experiment was analyzed using Refeyn DiscoverMP software (Version 2022 R1). Anti-PS solutions were prepared at different concentrations (0, 20, 50, 100, and 200 µg/ml) and incubated on the coated surfaces. Following interaction between the biomolecules and surfaces, the residual solution was collected and subjected to mass photometry analysis to determine the relative concentration of antibodies before and after surface immobilization.

### 4.7. ELISA assay to detect progesterone release

To evaluate the release of adsorbed progesterone from functionalized surfaces, glass coverslips coated with carboxyl-functionalized gold nanoparticles (GNPs) were incubated with 10 µM progesterone to allow adsorption. After adsorption, the coverslips were placed individually into wells of a 24-well plate and incubated with 150 µL of G-IVF™ Plus culture medium (Vitrolife) for defined time intervals ranging from 0 to 60 minutes, in triplicate. Each time point was assessed using separate samples to avoid cross-contamination.

Progesterone release into the culture medium was quantified using a competitive Progesterone ELISA kit (ALPCO, Cat. No. 11-PROHU-E01), according to the manufacturer’s instructions. Briefly, 25 µL of each sample (tested at 1:2 dilution), calibrator, and control were added in duplicate to the designated wells of the microplate, followed by the addition of 100 µL of 1× HRP-conjugated progesterone. The plate was incubated on a shaker at room temperature for 60 minutes. After three wash cycles with 300 µL of wash buffer per well, 150 µL of TMB substrate was added and incubated for 20 minutes. The reaction was stopped by adding 50 µL of stop solution, and the absorbance was measured at 450 nm using a microplate reader (BioTek). Progesterone concentrations were calculated by plotting optical densities against a standard curve generated using a 4-parameter logistic regression model.

### 4.8. Human semen preparation

Human semen samples were obtained from healthy donors as described in 4.1 above. Following liquefaction, a standard swim–up method was used to isolate motile sperm in prewarmed and gassed G–IVF^TM^ Plus media. Briefly, semen was overlayered 1:1 with medium and incubated at 37°C for 1 h. A carefully collected layer of motile sperm was evaluated for sperm count and motility. The remaining neat semen was used for sperm isolation using chemically-modified channel slides.

Relative yields are based on the percentage of progressively motile spermatozoa in the final preparation calculated using Mortimer’s method,^[56]^ according to the equation:

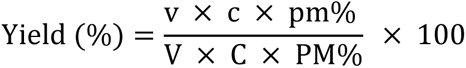

where v represents the volume of the sperm preparation (mL), c represents sperm concentration in the prepared sample (10^6^/mL), pm% represents progressive motility of the prepared sperm population (decimal), V represents initial volume of semen used for the procedure (mL), C represents sperm concentration in the semen sample (10^6^/mL), and PM% represents progressive motility in the semen sample (decimal).

### 4.9. Biocompatibility of the PPOx-coated surface

To evaluate the biocompatibility of the PPOx-coated surface, washed sperm at a concentration of 1 × 10^6^ in 100 µL G–IVF^TM^ Plus media were added to PPOx-coated glass coverslips in polystyrene tissue culture-grade petri dishes. PPOx-coated and uncoated standard glass coverslips were placed in 4–well plates (Thermo Scientific Nunc^TM^) and washed three times with copious sterile PBS for 15 min each time to remove any excess unreacted PPOx groups on the surface. The surface was then cleansed of potential microbial contaminants by incubation with 200 µg/mL streptomycin (Thermofisher Scientific), 200 U/mL penicillin (Thermofisher), and 500 ng/mL amphotericin B (Thermofisher) in sterile PBS for 2–3 h. Sperm were incubated at a density of 1 × 10^6^ per well on the coated or non– coated coverslip surfaces for 1 h and 5 h with G–IVF^TM^ Plus (Vitrolife) at 37°C and 5% CO_2_. Sperm incubated in polystyrene petri dishes (TCP) or on coverslips without coating were employed as controls.

### 4.10. Sperm attachment on the coated surface

The effect of incubating sperm in channel slides with different densities of surface immobilized anti–PS (10, 50, 100, 200, and 400 ug/mL) on sperm attachment was evaluated. 1 × 10^6^ swim-up sperm in 100 µL gassed G–IVF^TM^ Plus media was incubated on anti-PS coated surface for 30−60 min at 37°C and 5% CO_2_. Unattached sperm were removed from the surface and the number of attached sperm on the surface was quantified using light microscopy and the percentage of sperm attachment was calculated by dividing the number of attached sperms by the total number of sperm added to channel slides.

### 4.11. Unattached sperm quality

#### 4.11.1. Annexin V / PI Assay

The proportion of sperm labelled with the early–stage apoptotic marker Annexin V and cell viability marker PI was assessed in neat ejaculate, sperm obtained using conventional swim– up techniques, and unbound sperm populations collected following incubation with the immobilized anti–PS surface, using Annexin V / PI fluorescent labeling and flow cytometry. Annexin V–Alexa Fluor™ 488 (Thermofisher) was used to detect sperm in which translocation of PS from the inner to the outer leaflet of the plasma membrane has occurred (Annexin V^+^ sperm) indicating early stage apoptosis^[57]^, and PI was used to detect dead / late-stage apoptotic sperm (PI^+^ sperm). Briefly, harvested sperm were washed firstly with 1 mL PBS (350 x g, 5 min, 25°C), then samples were washed with 500 µL Annexin Binding Buffer (ABB) comprised of 50 mM HEPES, 700 mM NaCl, 12.5 mM CaCl_2_, pH 7.4 (Thermofisher). Sperm pellets were resuspended in ABB and aliquoted in 200 uL ABB at a final sperm density of 2 × 10^6^ in FACS tubes. Sperm samples were then gently mixed and 5 µL of Annexin V–FITC at a final concentration of 2.5 μg/mL and 4 µL PI at a final concentration of 2 µg/mL was added, followed by incubation for 5 min at room temperature. After incubation, sperm samples were washed in 500 µL ABB and then resuspended in ABB and analyzed within 10 min by FACSCanto II flow cytometer (BD Biosciences). Fluorescence signals were collected using appropriate filters (FITC: 530/30 nm; PI: 585/42 nm). Gating of stained populations was carried out using fluorescence intensity dot plots, allowing separation of live (Annexin V⁻/PI⁻), early apoptotic (Annexin V⁺/PI⁻), late apoptotic or necrotic (Annexin V⁺/PI⁺), and dead (Annexin V⁻/PI⁺) sperm populations. As a positive control to confirm the functionality of the Annexin V–FITC reagent, unprocessed spermatozoa were incubated at – 80 °C overnight to induce membrane damage and phosphatidylserine externalisation. These freeze-thawed spermatozoa reliably exhibited strong Annexin V binding (Supplementary Figure 5).

#### 4.11.2. Evaluation of sperm motility

Sperm motility was assessed according to WHO V guidelines.^[54a]^ Evaluation of sperm motility was measured on the CASA® semi-automatic semen analyzer (Microptic, Spain, Barcelona). 2 µL of sample was added to a Leja 10 slide and scanned using light microscopy (10x objective). A pre-set human count/motility program was used to calculate the proportion of progressive (STR >80%, where STR = straight linear velocity/average path velocity*100), non-progressive (STR <80%) and immotile sperm. Total motility was calculated by the proportion of progressive and non-progressive sperm. At least 200 sperm from at least five fields of view were evaluated.

#### 4.11.3. Evaluation of sperm DNA fragmentation by TUNEL

Sperm DNA fragmentation was evaluated using the TUNEL assay kit (Thermo Fisher) according to the manufacturer’s protocol, with minor modifications. Following attachment, unattached sperm were removed, and the attached spermatozoa were fixed by adding 1% (v/v) paraformaldehyde (Sigma Aldrich). The samples were incubated for 15 minutes at room temperature, followed by two washes with 5 mL PBS to remove excess paraformaldehyde.

Permeabilization was then performed by incubating the sperm samples in 0.1% Triton X-100 for 10 minutes at room temperature. After permeabilization, the sperm samples were washed twice with 1 mL Wash Buffer for 5 minutes. The TUNEL labelling reaction was carried out by incubating the sperm with 50 µL of labelling solution containing terminal deoxynucleotidyl transferase (TdT) for 2 hours at 37°C in the dark. Following the incubation, the sperm were washed twice with 1 mL Rinse Buffer for 5 minutes. To visualize the sperm nuclei, 100 µL of DAPI solution (1 µg/mL) was added to the samples, and the sperm were incubated for 10 minutes at room temperature in the dark. After DAPI staining, the samples were washed twice with 1 mL PBS to remove any excess dye. Fluorescence microscopy (Olympus IX73) was used to analyze TUNEL-positive sperm (green fluorescence) and DAPI-stained sperm (blue fluorescence).

#### 4.11.4. Evaluation of Sperm DNA Fragmentation

DNA fragmentation level was evaluated using Halosperm^®^ G2 kit (Halotech®) according to the manufacturer’s protocol. In brief, 100 µL agarose was placed in a 90 °C water bath for 30 min. It was then incubated at 37°C for at least 5 min before use. Following that, 50 μL of semen sample was gently mixed with the gel to avoid the bubble formation. Then, 8 μL of the mixed sample was pipetted onto an agarose–coated slide and covered with a coverslip. To solidify the gel, the slides were placed at 4°C for 5 minutes. The coverslip was then removed gently, and the slide was vertically submerged in solution 1 for 7 min at room temperature.

Then, the slide was immersed in solution 2 for 20 min and washed with distilled water for 5 min. The slide was dehydrated using 70% and 100% ethanol solutions for 2 min each. After air drying, the slides were incubated in solutions 3 and 4 for 7 min each step. To evaluate the percentage of sperm DNA fragmentation, the slides were imaged by a light microscope at x20 magnification. At least 100 sperm per slide were scanned to measure the ‘halo’ size using sperm class analyzer (SCA) software (Microptic L.S.). The size of the halo indicates the degree of sperm DNA fragmentation, with a large halo indicating intact DNA and a small or absent halo indicating fragmented DNA. The percentage of sperm with DNA fragmentation (%SDF) was calculated as the number of sperm with a small or absent halo as a percentage of total analyzed sperm.

### 4.12. Channel slide preparation

To compare the performance of the channel slide and swim-up selection techniques and to assess the effect of channel slide height on sperm selection and recovery, slides with channel heights of 0.1, 0.2, 0.4, 0.6, and 0.8 mm were evaluated. Semen samples were kept at 37°C for 30–60 min to liquefy. Channel slides were then prepared for use, by firstly loading the channels with an appropriate volume of G–IVF^TM^ Plus. The volume of G–IVF^TM^ Plus added was dependent on channel height, with volumes of 50, 100, 150 and 200 µL for channel heights of 0.2, 0.4, 0.6 and 0.8 mm respectively. The loading of the channels was performed slowly and gently, particularly in slides with low channel height (0.1 and 0.2 mm) to avoid trapping air inside the channel. Any trapped air acted to impede or completely prevent progressive sperm movement through the channel. Next, 60–100 µL of neat liquified semen was loaded into the inlet well, and the outlet well was filled with 60-100 µL G–IVF^TM^ Plus media. Slides were then incubated for 45–60 min at 37°C and 5% CO_2_ to allow progressively motile sperm to swim through the channel from the inlet well to the outlet well. Following incubation, the upper layer of the outlet well was carefully collected, and both sperm count and motility were assessed.

### 4.13. Statistical analysis

The data analysis was performed using GraphPad Prism GraphPad version 10.0.0 for Windows (GraphPad Software, Boston, Massachusetts USA, www.graphpad.com). The normality of continuous variables—including sperm motility, viability, apoptosis rate, and DNA fragmentation index—was assessed using the Shapiro–Wilk test to determine the appropriateness of parametric tests. Statistical significance between two groups was assessed using either an unpaired t-test or the non-parametric Mann–Whitney U test, as appropriate.

For comparisons involving three or more groups, a one-way ANOVA or a non-parametric Kruskal–Wallis test was applied. A significance level of P < 0.05 was considered indicative of a statistically significant difference between groups. Statistical significance was denoted as follows: *P<0.05, **P<0.01, ***P<0.001, and ****P<0.0001.

## Supporting Information

Supporting Information is available from the Wiley Online Library or from the author.

## Acknowledgements

This work was supported by project and fellowship grants from the National Health and Medical Research Council of Australia (APP1194466, APP1198172 and APP1145295).

## Data availability statement

The data that support the findings of this study are available from the corresponding author upon reasonable request.

