## Supplemental Figures 1-5 for "A surface-engineered microfluidic device for antibody-mediated negative selection of high-quality sperm for assisted reproduction"

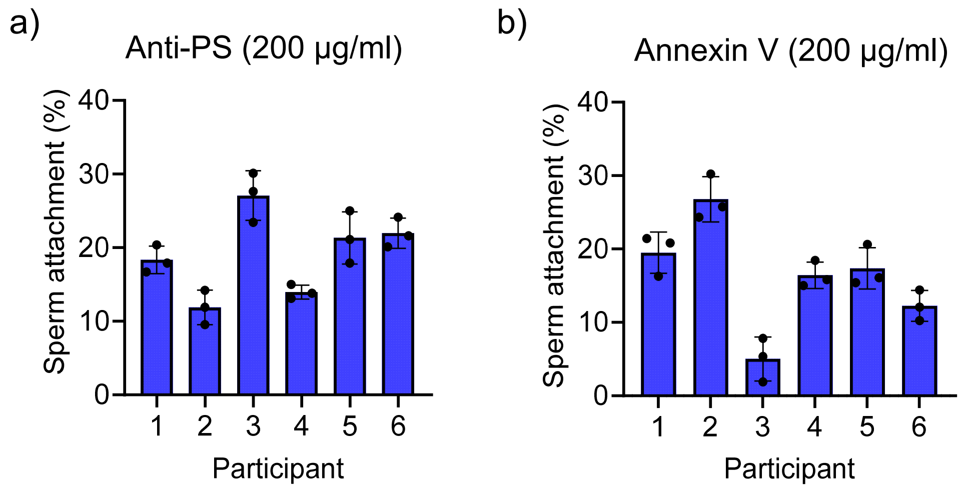


**Supplementary Figure 1**

The effect of incubating spermatozoa with Annexin V- and Anti-phosphatidyl serine (Anti-PS)-coated glass coverslips. (a) Percentage of spermatozoa adhesion to surface-immobilised Annexin V coated at a concentration of 200 µg/mL, and (b) Percentage of spermatozoa adhesion to surface-immobilised Anti-PS coated at a concentration of 200 µg/mL, utilizing swim-up spermatozoa samples obtained from a total of six individual normozoospermic men, with n=3 replicates of each.

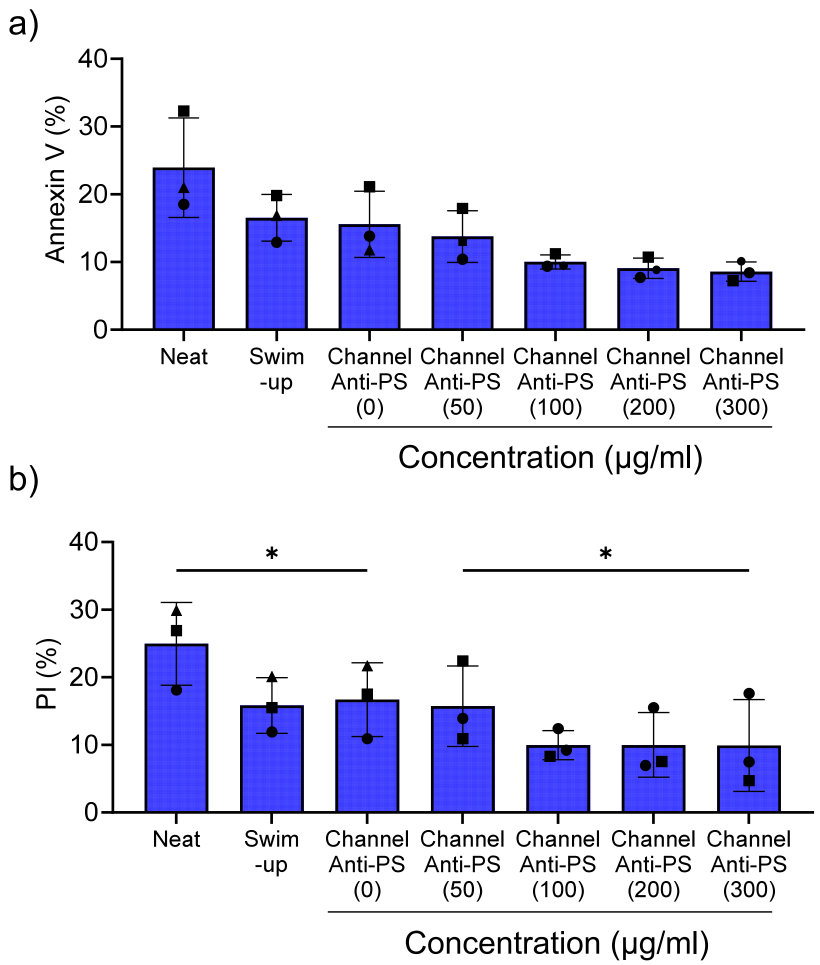


**Supplementary Figure 2**

Evaluation of the effect of concentration of anti-phosphatididyl serine (Anti-PS) on spermatozoa selection efficacy. Glass channel slides were coated with PPOx and then various concentrations of Anti-PS as described in the main text. Unselected spermatozoa (Neat), and spermatozoa processed by standard swim-up technique were included for comparison. (a, b) the proportion of recovered spermatozoa labeled with Annexin V and PI, indicating early and late-stage apoptotic populations respectively, after evaluation by flow cytometry. The results from each of 3 different spermatozoa preparations are shown using distinct symbols corresponding to each preparation. One-way ANOVA followed by post-hoc t-test was applied to evaluate the significance of differences between groups (*P≤0.05).

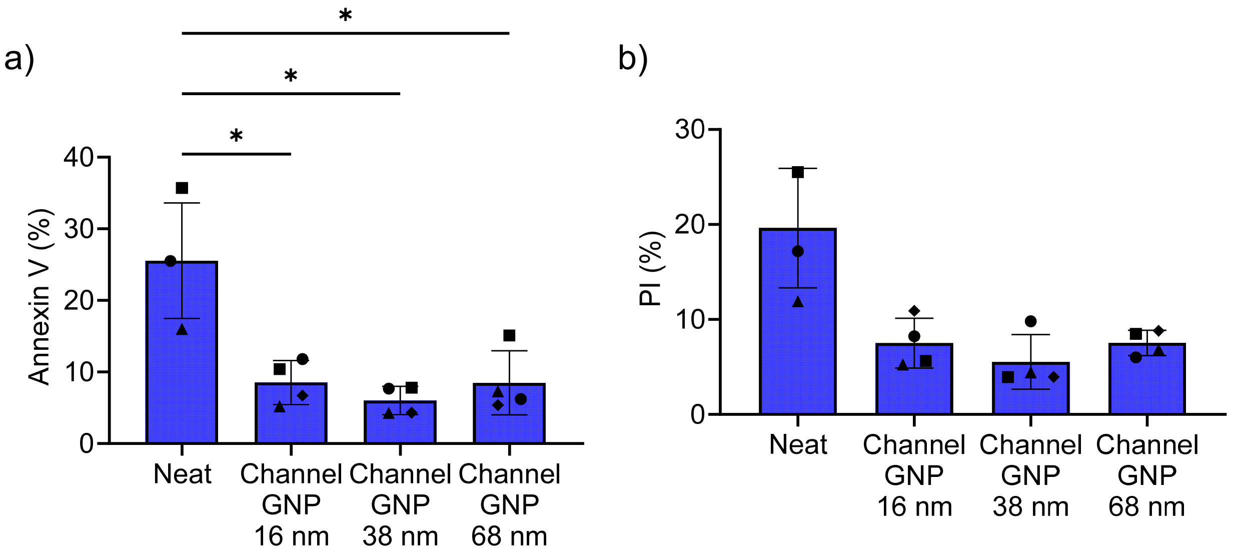


**Supplementary Figure 3**

Evaluation of the effect of gold nanoparticles (GNP) on spermatozoa selection. Glass channel slides were coated with PPOx and various sizes of GNP, then PPOx and anti-PS (200 ug/ml), as described in the main text. Unselected spermatozoa (Neat) were included for comparison. (a, b) the proportion of recovered spermatozoa labeled with Annexin V and PI, indicating early and late-stage apoptotic populations respectively, after evaluation by flow cytometry. The results from each of 3-4 different spermatozoa preparations are shown using distinct symbols corresponding to each preparation. One-way ANOVA followed by post-hoc t-test was applied to evaluate the significance of differences between groups (*P≤0.01).

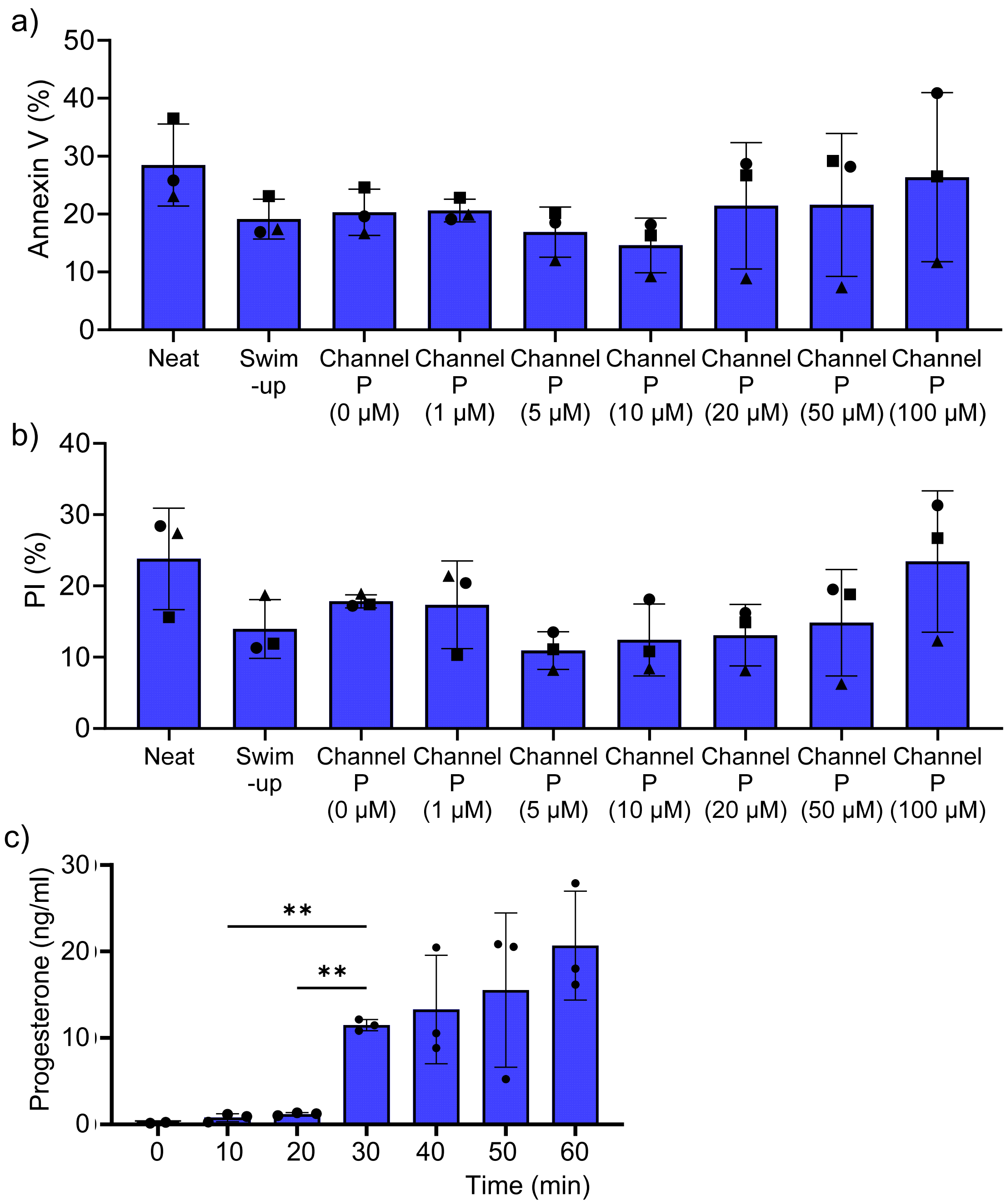


**Supplementary Figure 4**

Evaluation of the effect of progesterone (P) on spermatozoa. Progesterone at various concentrations was adsorbed onto glass channel slides coated with PPOx and GNP (68 nm) at the outlet chamber as described in the main text. Unselected spermatozoa (Neat), and spermatozoa processed by standard swim-up technique were included for comparison. (a, b) the proportion of recovered spermatozoa labeled with Annexin V and PI, indicating early and late-stage apoptotic populations respectively, after evaluation by flow cytometry. The results from each of 3 different spermatozoa preparations are shown using different symbols for each sperm sample. No significant differences were seen when one-way ANOVA followed by post-hoc t-test was applied to evaluate differences between groups. (c) Time-dependent release of progesterone from carboxyl-functionalized gold nanoparticle (GNP)-coated coverslips into G-IVF™ culture medium, quantified using a competitive ELISA assay. Coverslips were pre-incubated with 10 µM progesterone and then placed in 150 µL of G-IVF medium for 0, 15, 30, 45, or 60 minutes. At each time point, the medium was collected from independent samples and analyzed for progesterone concentration. Data are presented as mean ± standard deviation (SD) and (**P≤0.01).

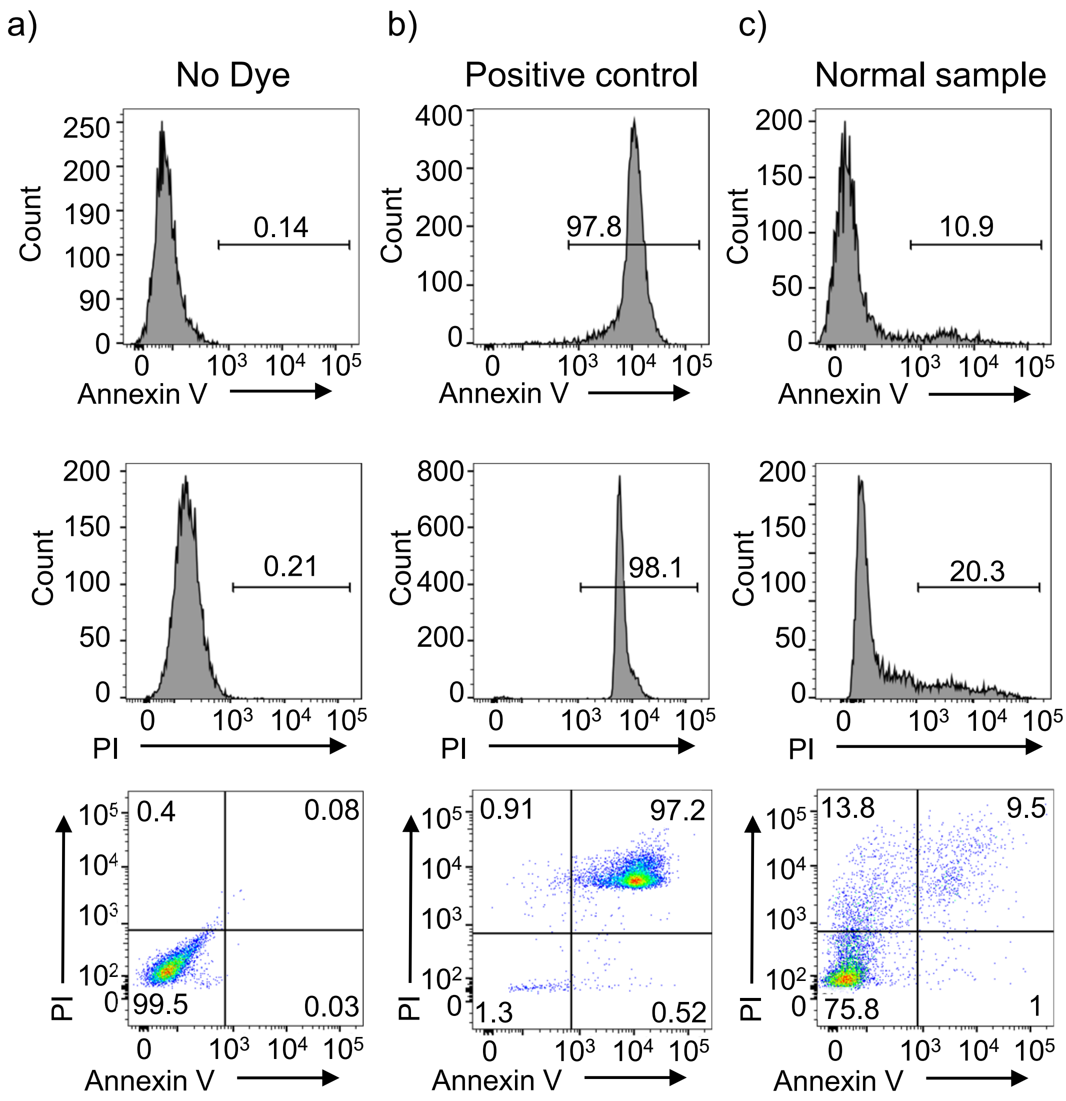


**Supplementary Figure 5**

Flow cytometry analysis of spermatozoa stained with Annexin V–FITC and propidium iodide (PI). (a) Control plot demonstrating flow cytometry gating. (b) Positive control: spermatozoa stored at –80 °C overnight to induce membrane damage and phosphatidylserine externalisation. (c) Freshly collected, untreated spermatozoa stained with Annexin V–FITC and PI. (a-c) From top to bottom, histograms represent Annexin V–positive population, PI–positive population, and the dot plot of dual staining (Annexin V vs PI). Numbers in quadrants are %sperm.
